# Hippo pathway effectors YAP1/TAZ induce a EWS-FLI1-opposing gene signature and associate with disease progression in Ewing Sarcoma

**DOI:** 10.1101/589648

**Authors:** Pablo Rodríguez-Núñez, Laura Romero-Pérez, Ana T. Amaral, Pilar Puerto-Camacho, Carmen Jordán, David Marcilla, Thomas G. P. Grunewald, Enrique de Alava, Juan Díaz-Martín

**Author notes:** Corresponding authors: Enrique de Álava,; Juan Díaz-Martín,; Laura Romero-Pérez. Equal contribution (Pablo Rodríguez-Núñez, Laura Romero-Pérez, Juan Díaz-Martín).

## Abstract

YAP1 and TAZ (WWTR1) oncoproteins are the final transducers of Hippo tumor suppressor pathway. Deregulation of the pathway leads to YAP1/TAZ activation fostering tumorigenesis in multiple malignant tumor types, including sarcoma. However, oncogenic mutations within the core components of the Hippo pathway are uncommon. Ewing Sarcoma (EwS), a pediatric cancer with low mutation rate, is characterized by a canonical fusion involving *EWSR1* gene, and *FLI1* as the most common partner. The fusion protein is a potent driver of oncogenesis but secondary alterations are scarce, and little is known about other biological factors that determine the risk of relapse or progression. We have observed YAP1/TAZ expression and transcriptional activity in EwS cell lines. Analyses of 55 primary human EwS samples revealed that high YAP1/TAZ expression was associated with progression of the disease and predicted poorer outcome.

We did not observe recurrent SNV or copy number gains/losses in Hippo pathway-related loci. However, differential CpG methylation of *RASSF1* locus -a regulator of Hippo pathway- was observed in EwS cell lines compared with mesenchymal stem cells, the putative cell of origin of EwS. Hypermethylation of *RASSF1* correlated with the transcriptional silencing of the tumor suppressor isoform *RASFF1A*, and transcriptional activation of the protumorigenic isoform *RASSF1C* promoting YAP1/TAZ activation. Knockdown of YAP1/TAZ decreased proliferation and invasion abilities of EwS cells, and revealed that YAP1/TAZ transcription activity is inversely correlated with the EWS-FLI1 transcriptional signature. This transcriptional antagonism could be partly explained by EWS-FLI1-mediated transcriptional repression of TAZ. Thus, YAP1/TAZ may override the transcriptional program induced by the fusion protein, contributing to the phenotypic plasticity determined by dynamic fluctuation of the fusion protein, a recently proposed model for disease dissemination in EwS.

## INTRODUCTION

Ewing Sarcoma (EwS) represents the second most common primary malignant bone tumor in children and young adults [1]. Owing to multimodal treatment concepts, 2/3 of the patients with localized disease achieve sustained remission but approximately 30 % relapse. Patients at relapse or with advanced disease have limited chance to survive with a three-year event free survival of less than 25% [2, 3]. While clinical prognostic markers such the presence of metastases or tumor volume are established, little is known about the biological factors determining the risk of progression, thus precluding risk-adapted therapeutic approaches. EwS was the first solid malignancy defined by the presence of tumor-specific EWSR1-ETS gene fusions [4], mainly EWSR1-FLI1 translocations, which are considered the main driver of the disease, but fusion type itself does not have any impact on disease progression [5]. As in most developmental cancers, additional recurrent mutations are scarce. The most common somatic mutations have been detected in *STAG2*, *CDKN2A and TP53*, associated with poor prognosis [6, 7]. Copy number variation studies by the PROVABES consortium using samples derived from the EURO-E.W.I.N.G.99 (EE99) and EWING 2008 trials showed that chromosome 1q gain and possibly chromosome 16q loss define patients with poor clinical outcome (Díaz-Martín et al, unpublished data), supporting previous retrospective studies [7, 8]. However, these secondary alterations occur with a frequency which does not account for the large proportion of patient who relapses.

The Hippo tumor suppressor pathway plays a critical role in tissue and organ size regulation by restraining cell proliferation and apoptosis under homeostatic conditions [9]. Central to Hippo pathway is a conserved cascade of adaptor proteins and inhibitory kinases that regulate the activity of the oncoproteins YAP1 and TAZ, the final effectors of this pathway in mammals. YAP1/TAZ do not directly bind DNA, but act as transcriptional coactivators of target genes involved in cell proliferation and survival through their interaction with transcriptional regulators such as TEAD factors [10]. The role of YAP1 and TAZ as important drivers in tumorigenesis has been extensively reported in carcinomas, and they also contribute to malignancies of mesenchymal origin [11–13]. In fact, given its key function in developmental processes, an important role has been inferred for Hippo signaling in pediatric cancer [14]. Despite this, somatic or germline mutations in Hippo pathway genes are uncommon, in comparison to other well-defined signaling pathways that are commonly disrupted in cancer [13, 15]. Since secondary genetic alterations are scarce in EwS, and given the established role of YAP1 and TAZ in cancer without engaging mutation, we aimed to explore the contribution of these factors to oncogenesis in Ewing sarcoma. Herein we evaluated a series of 55 EwS patients by immunohistochemistry (IHC) for expression/activation of YAP1 and TAZ. We observed a significant association of YAP1/TAZ nuclear expression and disease progression, as well as a potential mechanism of dysregulation involving epigenetic regulation of *RASSF1* locus. Moreover, we demonstrated an interesting interplay between TAZ/YAP1 function with the fusion protein, which fits into a recent model concept for metastatic spreading in EwS based on fluctuations of the expression of the fusion protein [16].

## MATERIALS AND METHODS

### Tumor samples

In this study we analyzed 88 formalin-fixed paraffin-embedded (FFPE) samples from 68 Ewing sarcoma patients (55 samples corresponding to primary tumor). We also analyzed a subset of 21 frozen samples from the same series. Clinical diagnosis of all the samples was performed according to the World Health Organization (WHO) classification [17], performing fluorescence in situ hybridization (FISH) to assess the presence of EwS translocation in tissue sections, which validates the immunohistochemical diagnosis. The only selection criteria were the availability of pathological data and tissue for tissue microarray (TMA) construction. Medical records were retrospectively reviewed and clinicopathologic information for 55 patients with primary tumor material were retrieved for further analyses (summarized in **Table 1**). Tissue samples were obtained from the HUVR-IBiS Biobank (Universitary Hospital Virgen del Rocio-Institute of Biomedicine of Seville Biobank. Andalusian Public Health System Biobank). This study was performed following the standard Spanish ethical regulations and it was approved by the corresponding ethics committee of the Hospital Virgen del Rocío de Sevilla and the Fundación Pública Andaluza para la Gestión de la Investigación en Salud de Sevilla (FISEVI), Spain. Written informed consent was obtained from all patients and all clinical analyses were conducted in accordance with the principles of the Helsinki Declaration.

**Table 1.** Clinical and pathologic findings according to YAP1/TAZ nuclear expression in primary EwS specimens (n=55)

| Characteristics | Analyzable | TAZ/YAP1 expression (IHC) |  | p |
| --- | --- | --- | --- | --- |
|  |  | Negative/Weak positive | Strong positive |  |
| **Age (years) mean** | **55** | 20,72 (±2,062) | 20,93 (±3,196) | 0.9088 |
| **Location** | **42** |  |  | 0.4945 |
| Bone | 28 (66,67%) | 19 (67,86%) | 9 (32,14%) |  |
| Soft tissue | 14 (33,33%) | 8 (57,14%) | 6 (42,86%) |  |
| **Progression** | **50** |  |  | **0.0054** |
| No | 27 (54%) | 21 (77,78%) | 6 (22,22%) |  |
| Yes | 23 (46%) | 9 (39,13%) | 14 (60,87%) |  |

### TMA construction and Immunohistochemistry

Representative tumor areas of EwS samples were selected on H&E-stained sections and two 1-mm diameter tissue cores were obtained from each specimen to set up 4 different TMAs. IHC was carried out on TMA sections using the Envision method (Dako, CA, USA) with a step of heat-induced antigen retrieval and using a primary antibody against YAP1 and TAZ (**Suppl. Table 1**). IHC staining was separately evaluated by two pathologists. YAP1/TAZ expression was evaluated for nuclear staining, thus focusing in their transcriptional activity. Tissue was given a score which resulted of multiplying the nuclear staining intensity from 0 (no staining) to 3 (strong staining), by the extension based on the percentage of positive cells (from 0 to 3). Samples were grouped as negative or weak positive (score 0-2), and strong positive (3-9).

### Cell lines

EwS cell lines SKNMC, TTC-466, TC32, A4573, A673, CADO-ES, RD-ES, RM82, SKES1, STAET10, TC71 and WE68, were obtained from the EuroBoNet cell line panel [18]. MDA-MB-231, MCF7, RH30, SAOS2 and PC3 cell lines were purchased from ATCC. Primary human bone marrow mesenchymal cells (hMSC), immortalized with telomerase reverse transcriptase were provided by D. Campana [19]. Each cell line was grown in its corresponding culture medium (DMEM, RPMI, EMEM or McCoýs) supplemented with 10-15% of FBS (Fetal Bovine Serum) and 1% penicillin-streptomycin. We also used EwS cell line A673 engineered to express a doxycycline-inducible shRNA against the EWS-FLI1 fusion protein [20]. Cells were grown in DMEM supplemented with 10% fetal bovine serum, 100 μg/ml Zeocin (InvivoGen, ant-zn-5p) and 5 μg/ml Blasticidin (InvivoGen, ant-bl-10p). For the EWS-FLI1 shRNA induction, 1μg/ml doxycycline (Sigma, D9891) was added to the media for 48 hours.

### Western Blotting

Western Blotting was performed to analyze the expression of different proteins by using primary antibodies detecting: YAP, SRC, Phospho-SRC, TAZ, CYR61, CTGF and Calnexin/GAPDH as endogenous controls. Antibodies details are provided in Supplementary Table 2. Blot detection was carried out by using Clarity western peroxide reagent (Bio-Rad Clarity western ECL substrate) and visualized by digital imaging in a Chemidoc Touch Imaging System (Bio-Rad).

### Nucleus and cytoplasm subcellular fractionation

Cells (∼9×106) were washed twice with ice-cold 1× PBS. Citoplasmic lysis buffer (HEPES pH 7,9, KCl 10 mM, EDTA 0,1 mM, EGTA 0,1 mM, PMSF 0,5 mM, DTT 1mM, NaF 1mM, Na3VO4 1mM, PIC 1x, Nonidet NP40×0,625%) is added and attached cells were scraped off with a cell scraper. Cells were centrifugated for 2 minutes at 13000 RPM. Citoplasmic fraction-containing supernatant is placed in a different eppendorf and the nuclear fraction-containing pellet is resuspended in a Nuclear lysis buffer (HEPES pH 7,9 20 mM, NaCl 0,4M, EDTA 1 mM, EGTA 1 mM, PMSF 1mM, DTT 1mM, NaF 1mM, Na3VO4 1mM, PIC 1x) at 4°C for 15 minutes with vortexing for 30 seconds every 5 minutes. After centrifugation for 5’ at 14000 RPM, nuclear fraction-containing supernatant was isolated.

### Luciferase assays

Cells were seeded in 24 well plates 24h before transfection. The established YAP1/TAZ-TEAD responsive reporter 8xGTII–lux was a gift from Stefano Piccolo (Addgene plasmid # 34615) [21]. As negative control reporter we used pTNT-min (provided by Mark Bond [22]), lacking the TEAD elements. Luciferase reporters (400 ng/well) were transfected with Lipofectamine LTX with Plus Reagent (Thermo Fisher Scientific) together with pRL-TK Renilla (50 ng/well) to normalize for transfection efficiency. Cells were collected 24h after DNA transfection. Cell lysates were analysed using the Dual-Luciferase Reporter Assay System (Promega, #E1910). Luminiscence was measured using a TECAN infinite M200-PRO plate reader (Tecan, Männedorf, Switzerland).

### siRNAs

Silencing of TAZ and YAP1 was performed by using 10nM Silencer Select siRNAs (Thermo Fisher Scientific), whose sequences are detailed in **Suppl. Table 1**. Viromer Blue was used as a transfection reagent according to the manufactureŕs conditions (Lipocalyx, Germany). We evaluated the eficiency of two different pairs of siRNAs targeting YAP1 and TAZ, and we confirmed similar results for both (**Suppl. Fig. 4**). We used a negative control designed against a non-human coding sequence (siC) and a positive control designed against GAPDH (siG).

### Drugs

Dasatinib and pitavastatin were purchased from Selleckchem (Houston, TX, USA). Stock solutions of both compounds were prepared in dimethyl sulfoxide (DMSO) and diluted to final concentration in the culture medium. Controls were treated with DMSO at the same final concentration. The DMSO never exceeded 1:1000 (v/v) of total incubation volume and did not show any toxic effects on EwS cells.

### Cell proliferation assay

Cell proliferation was evaluated using the ATP-lite 1 step Luminescence Assay System (Perkin Elmer). Cells were seeded in gelatin pre-treated 96 wells plates, avoiding marging areas, for at least 24 h. Culture medium was removed and ATP-lite solution was added. Luminiscence was measured using a TECAN infinite M200-PRO plate reader (Tecan, Männedorf, Switzerland).

### Migration assay

Migration assay was performed as described previously [23]. Cells were grown on 6 wells plates to 85-90% confluence and a wound was made by scratching the monolayer of the cells. Pictures of the same selected area were taken after the times indicated and the percentage of wound healing was calculated using the imageJ software.

### Invasion assay

Invasion assay was carried out on modified Boyden chambers (8 µm pore filters) from Cultrex (Trevigen). These chambers were coated with BME (Cultrex Basement Membrane Extract), a natural extracellular matrix hydrogel. It was diluted at 0.2X in coating buffer and added to the inserts 24 h before cells were seeded. Cells were cultured in medium without serum for at least 24 h before the assay. Afterwards, a volume of 100 µl of serum-free medium containing 1×10^5^ cells was deposited on the top of the chamber. Serum was used as chemoattractant in the lower part of the chamber. After 48h at the incubator at 37C, both serum-free medium from the top and pure serum from the lower part of the chamber were removed. The top of the membrane was carefully cleaned and the bottom retaining invading cells was washed with PBS. Cells were fixed to the membrane with methanol 100% for 5 minutes at −20°C. After PBS washing, cells were stained with DAPI and membranes were mounted on a slide to be observed under a fluorescence microscope (Olympus BX-61). Cell nucleus of the respective conditions were counted and compared with controls.

### Transcriptome analysis

SK-N-MC cells were transfected with control or a combination of YAP1/TAZ siRNAs for 72h. Whole transcript expression analysis was conducted in four biological replicates of each sample. RNA was amplified and labeled using the GeneChip® WT PLUS Reagent Kit (Thermo Fisher Scientific, Inc.). Amplification was performed with 100 ng of total RNA input following procedures described in the WT PLUS Reagent Kit user manual. The amplified cDNA was quantified, fragmented, and labeled in preparation for hybridization to GeneChip® Human Transcriptome 2.0 Array (Thermo Fisher Scientific, Inc.) using 5.5 μg of single-stranded cDNA product and following protocols outlined in the user manual. Washing, staining (GeneChip® Fluidics Station 450, Thermo Fisher Scientific, Inc.), and scanning (GeneChip® Scanner 3000, Thermo Fisher Scientific, Inc.) were performed following protocols outlined in the user manual for cartridge arrays. The fluorescence signals scanned as DAT files were transformed to CEL files via the AGCC software (Thermo Fisher Scientific, Inc.). The GeneChip® Command Console software (Thermo Fisher Scientific, Inc.) pretreated CEL files through robust multichip analysis algorithm to obtain CHP files. Next, CHP files were analyzed by Transcriptome Analysis Console (TAC) 4.0 software (Thermo Fisher Scientific, Inc.) which performs statistical analysis and provides a list of differentially expressed genes. Gene set enrichment analysis (GSEA v3.0) was performed to identify targets of YAP1/TAZ that are over-represented in previous defined gene sets [24, 25].

### Genome-wide copy number analysis

FFPE samples were sliced into 10-μm sections and gDNA was extracted using the QIAamp DNA FFPE Tissue Kit (Qiagen, Sussex, United Kingdom). DNA concentration was determined using the Quant-iT™ PicoGreenR dsDNA Assay Kit (Thermo Fisher Scientific UK Ltd., Paisley, United Kingdom). Genome-wide copy number analysis was performed using the OncoScan FFPE Assay Kit (Affymetrix, Santa Clara, CA, USA) according to the manufacturer’s recommendations. Nexus Express for OncoScan 3 software (BioDiscovery, Hawthorne CA, USA) was used to estimate copy numbers. Significance testing for aberrant copy number (STAC) method was conducted to evaluate the significance of DNA copy number aberrations across the tumor series.

### Methylation array

Methylation data were generated as described in Puerto-Camacho et al. (2018) [26]. Data analyses (GSE118872) were performed using the Bioconductor lumi package [27].

### Statistical analysis

Correlation between inmunohistochemical YAP1/TAZ expression and clinicopathological characteristics was assessed by chi-squared test for the categorical variables (summarized with percentages). Mann–Whitney test was used for the analysis of differences of the continuous variable age (summarized with means and standard errors). EwS-specific survival was defined as the time from surgery to the time of death from EwS with deaths from other causes being censored, whereas in time to relapse analysis, the end point was EwS recurrence, either local or distant. Survival curves were estimated using the Kaplan-Meier method, and the differences in survival were evaluated using the long-rank test. Cox’s proportional hazards modeling of parameters potentially related to survival were conducted to calculate hazard ratios (HR), in both univariate and multivariate analyses. All these statistical analyses were performed using SPSS version 20 (SPSS Inc., Chicago, IL, USA) and JMP 10 statistical software (SAS Institute Inc., Cary, NC, US). p< 0.05 was considered statistically significant.

Statistical analysis of in vitro functional assays was performed by using SPSS 20.0 (SPSS Inc., Chicago, IL, USA), and represented by graph tools from Excel (Microsoft Office v.10) or Origin Pro 9.0 software.

## RESULTS

### YAP1/TAZ are expressed in EwS cell lines and tumor specimens, and are associated with the presence of metastasis and poor prognosis

First, we examined YAP1/TAZ expression by WB in 13 EwS cell lines with different pathognomonic gene fusions (**Fig. 1A**). We observed heterogeneous expression of both proteins across the cell line panel. Some of the EwS cell lines showed YAP1/TAZ expression comparable to cell lines in which a relevant role has been described for these factors (i.e. MDA-MB-231, a triple negative breast cancer cell line with NF2- mutations leading to activation of TAZ/YAP1) [**28**]. TAZ/YAP1 expression was also detected in human mesenchymal stem cells (hMSC) derived from bone marrow, a proposed cell of origin of EwS. Importantly, nuclear expression was observed by subcellular fractionation and immunofluorescence (**Fig. 1B-C**, **Suppl. Fig. 1**), suggesting functional transcriptional activity which was confirmed with luciferase reporter assays (**Fig. 1D**).

**Figure 1.**
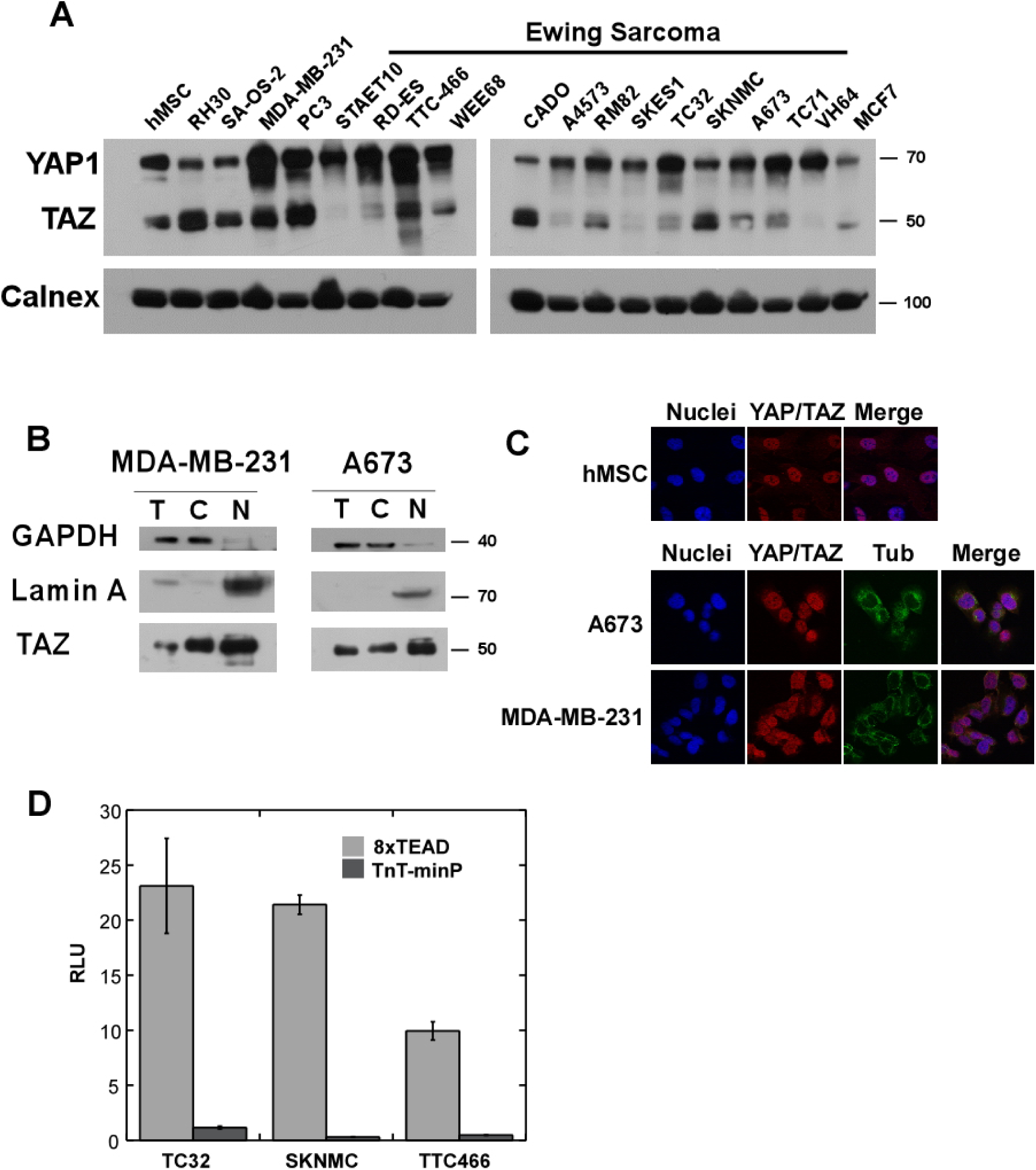
YAP1 and TAZ are expressed and active in EwS cell lines. (**A**) Western blot using a monoclonal antibody recognizing total levels of YAP1 and TAZ proteins in a panel of 13 EwS cell lines. Basal and luminal breast cancer (MDA-MB-231, MCF-7), prostate cancer (PC3), osteosarcoma (SA-OS-2), rhabdomyosarcoma (RH30) and human mesenchymal stem cells (hMSC) were included in the assay.(**B**) Nucleus and cytoplasm subcellular lysates were assessed by WB (T, Total extract; N, nucleus; C, cytoplasm). (**C**) Immunofluorescence microscopy with the indicated antibodies (60X). (**D**) YAP1/TAZ-TEAD dependent transcriptional activity in EwS cell lines was evaluated with luciferase reporter constructs containing sequences with or without TEAD elements (8xTEAD and TnT-minP constructs respectively). RLU (Relative luminescence units) was normalized to renilla luciferase values. Data are represented as mean ±SE of 3 biological replicates.

To test whether YAP1/TAZ abundance was associated with clinical variables in EwS, we analyzed their expression by IHC in a retrospective series of 55 primary tumors (**Table 1**). YAP1/TAZ strong expressing tumor cells exhibited intense nuclear staining with a variable signal in the cytoplasm (**Fig. 2A**). YAP1/TAZ expression was also observed in endothelial cells in negative samples providing an internal positive control for the IHC determination (**Fig. 2B**). YAP1/TAZ strong expression was associated to disease progression (chi-square test, p<0.0054), whereas no significant association was observed with age at surgery or location (**Table 1**). We also observed increased YAP1/TAZ positivity in metastatic or relapsed tumors in 11 patients with paired samples (**Fig. 2C-H, 2I**, paired t-test, p = 0.0204). Additional non-paired metastatic or relapsed tumor samples showed preferential strong expression as well (**Fig. 2J**, Fisher’s exact test, p = 0.006).

**Figure 2.**
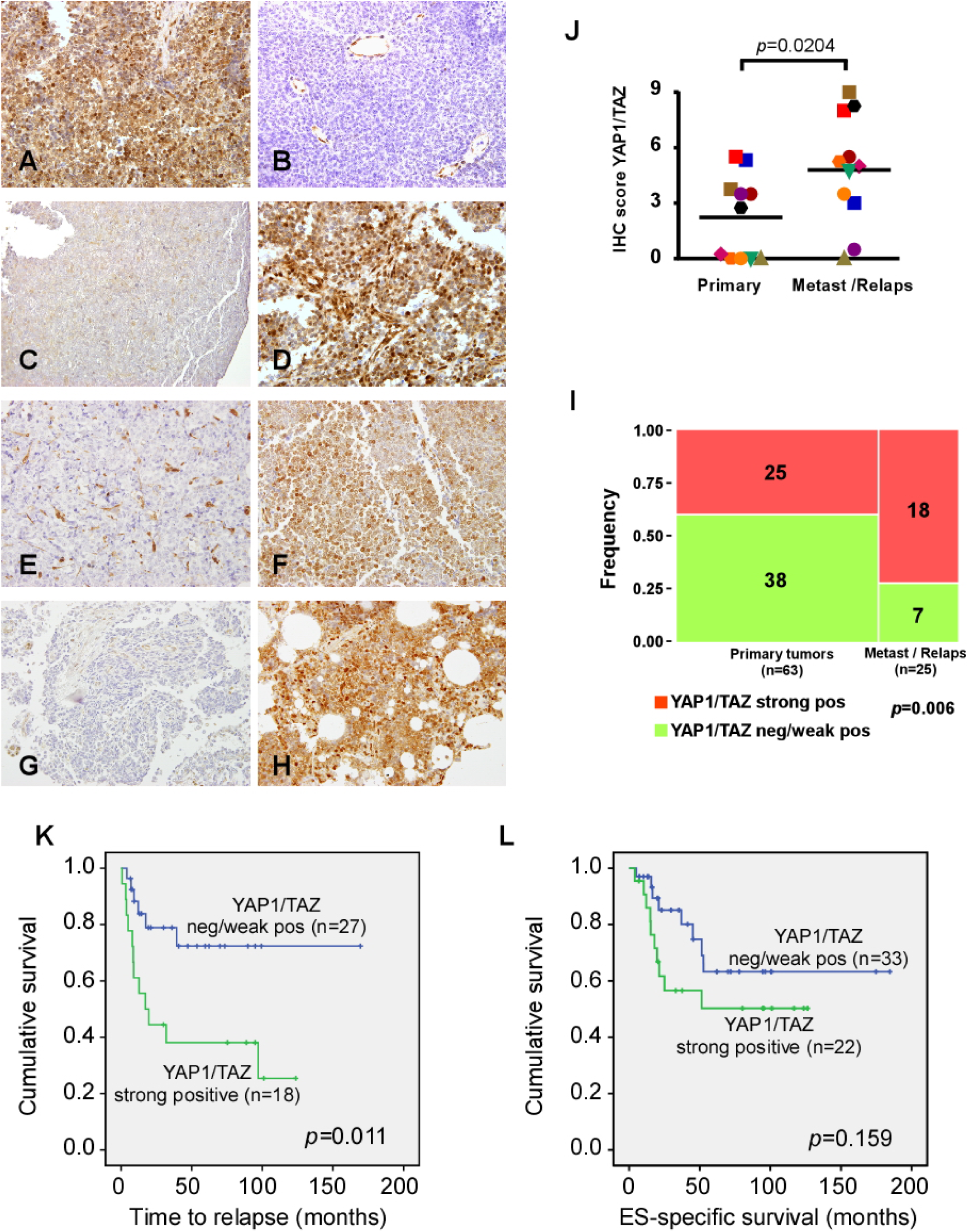
YAP1/TAZ expression associates with disease progression. (**A**) Representative image for YAP1/TAZ strong positive expression in a primary EwS tumor (40x). (**B**) Staining of endothelial cells can be observed in a negative tumor specimen. (**C-H**) Immunostaining for YAP1/TAZ in primary tumors (left) and matched metastasis (right) of the same patients (40x). (**I**) Comparison of YAP1/TAZ immunostaining in 11 matched biopsies. (**J**) Distribution of samples in each tumor category (primary vs metastasis or relapse) according to YAP1/TAZ staining score. The number of samples is indicated on the bars. (**K**, **L**) Kaplan-Meier survival curves for YAP1/TAZ protein expression in EwS patients grouped as negative/weak positive vs strong positive staining.

We retrieved follow-up data for the EwS patients with primary tumor biopsies to evaluate prognosis (median duration of follow-up of 35.23 months), but only 45 had known relapse date (median duration of follow-up of 41.43 months). YAP1/TAZ expression influenced significantly the time to relapse which was shorter in strong positive patients than in weak/negative patients (mean, 127.4 vs. 50.66 months, p=0.011, **Fig. 2K**). Similarly, Kaplan-Meier estimates of EwS specific survival were shorter (but not significant) for the YAP1/TAZ strong positive group compared with YAP1/TAZ weak/negative group (mean, 129.32 vs. 73.61 months, p=0.159, **Fig. 2L**). Accordingly, Cox regression univariate analyses determined that YAP1/TAZ strong expression was significantly correlated with the time to relapse but not with EwS specific survival, with the unadjusted hazard ratio (HR) being 3.354 (p=0.016) and 1.928 (p=0.167) respectively (**Table 2**). A significant correlation with survival and time to relapse was also observed for metastasis (**Table 2**). These variables were all included simultaneously, to assess the independent prognostic significance based on multivariate analysis. The adjusted HR of YAP1/TAZ strong expression for relapse did not reach significant confidence regarding time to relapse, after controlling the Coxˈs regression model for the effects of age, tumor location and metastasis. However, a roughly significant HR for YAP1/TAZ was obtained in the multivariant analysis (**Table 2**).

**Table 2.** Prognostic value of YAP1/TAZ IHC expression in relation to other clinical variables.

| Factors | Time to relapse |  |  |  | EwS-specific survival |  |  |  |
| --- | --- | --- | --- | --- | --- | --- | --- | --- |
|  | Unadjusted HR (95%CI) | p | Adjusted HR (95%CI) | p | Unadjusted HR (95%CI) | p | Adjusted HR (95%CI) | p |
| **TAZ/YAP1 (strong vs. negative/weak)** | 3,354 (1,253-8,974) | **0.016** | 1,579 (0,287-8,676) | 0.599 | 1,928 (0,761-4,886) | 0.167 | 5,703 (1,004-32,400) | **0.049** |
| **Metastasis** | 14,895 (4,675-47,455) | **<0,001** | 70,369 (4,980-994,326) | **0.002** | 11,318 (3,211-39,895) | **<0,001** | 77,954 (7,332-828,754) | **<0,001** |
| **Age** | 0,967 (0,916-,1022) | 0.234 | 1,034 (0,924-1,159) | 0.557 | 0,970 (0,925-1,017) | 0.208 | 0,971 (0,888-1,062) | 0.523 |
| **Location (bone vs soft tissue)** | 2,491 (0,667-9,294) | 0.174 | 2,612 (0,468-14,594) | 0.274 | 1,066 (0,326-3,483) | 0.916 | 2,640 (0,478 -14,570) | 0.265 |

### Activation of YAP1 /TAZ in Ewing Sarcoma

We tried to determine the mechanisms that contribute to YAP1/TAZ activation in EwS. To do so, we interrogated public datasets for somatic mutations in the Hippo pathway-related genes, but we did not find any recurrent SNV (**Supp. Fig. 2**). Next, we analyzed copy number alterations in a series of 24 EwS by SNP arrays (**Fig. 3**). Gross chromosomal alterations were similar to previous reports, i.e. gains of whole chromosomes 8 and 12 [**7**]. Copy number gain in *WWTR1* locus, with complete gain of chromosome 3 was detected in a single case. Gain at *YAP1* locus was detected in another case with an almost tetraploid genotype. None of the two cases showed incremented mRNA expression associated to the copy number event. Regarding the core regulatory kinases of the Hippo pathway and other negative regulators of YAP1/TAZ function, no significant copy-loss events were observed (**Fig. 3**). Focal copy number aberration events in Hippo-related loci were also precluded after inspecting the data with the STAC algorithm (**Suppl. Table 2**). Similarly, Hippo-related loci were unaffected in a retrospective series of 165 cases of EwS, which was analyzed within the PROVABES consortium for validation of biomarkers in EwS (www.medizin.uni-muenster.de/provabes/network, Díaz-Martín J., unpublished data).

**Fig. 3.**
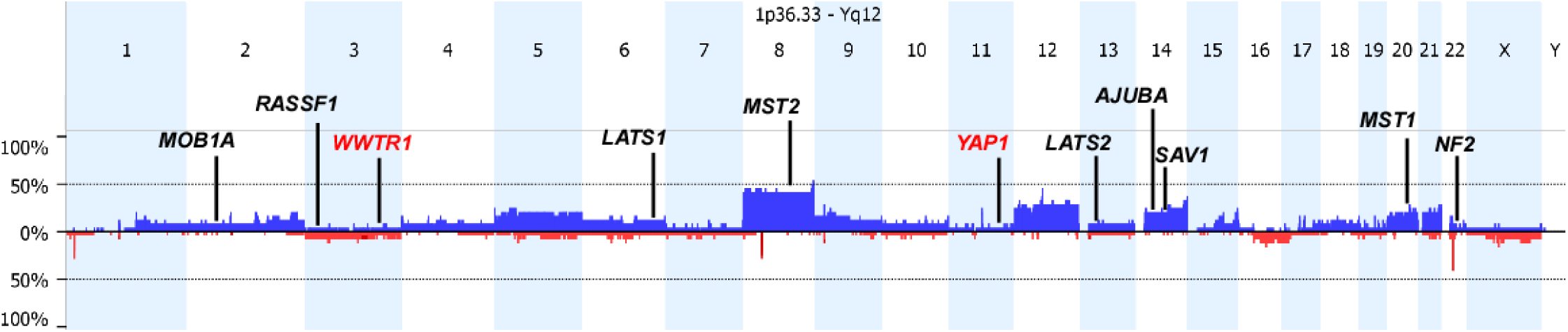
Summary of copy number aberrations detected in 24 EwS samples. Frequencies of copy number gain (above axis, blue) and copy number loss (below axis, red) across the human genome. Hippo-related loci are indicated: Tumor suppressor genes such us core kinases of the pathway are marked in black, and oncogenes *WWTR1* and *YAP1* in red.

Deregulation of Hippo pathway leading to YAP1/TAZ activation could be the consequence of epigenetic silencing of tumor suppressor genes through DNA hypermethylation [11, 15, 29]. We inspected previous results of the group comparing CpG methylation in EwS cell lines versus human mesenchymal stem cells (hMSC) from EwS patients and healthy donors (**GSE118872**) [26]. Among the differentially methylated genes, we found that one of the negative regulators of YAP1/TAZ, *RASSF1*, was hypermethylated in EwS cells. No other Hippo-related loci showed differential methylation (**Fig. 4A**). Hypermethylation of *RASSF1* accounts for silencing of *RASSF1A* transcript expression, but promotes switching to an alternative gene promoter driving the expression of the isoform *RASSF1C*. *RASSF1A* contributes to Hippo pathway-mediated repression of YAP1/TAZ, whereas *RASSF1C* promotes Src family kinases (SFKs)-mediated activation of YAP1 [30]. We confirmed expression of the alternate isoform *RASSF1C* in EwS cell lines, whereas *RASSF1A* expression was absent or reduced (with the exception of STAET-10 and TC-32 cell lines) compared with hMSC (**Fig. 4B**). Moreover, expression of YAP1/TAZ target genes positively correlated with *RASSF1C* expression in the cell line panel, as well as in EwS tumor specimens (**Fig. 4C, D**). Interestingly, TAZ but not YAP1 seems to be transcriptionally regulated since CTGF expression correlate with TAZ mRNA expression (**Fig. 4D**). Correlation of TAZ mRNA levels with Hippo target genes was also observed in larger EwS series in public repository expression data (**Suppl. Fig. 3**).

**Fig. 4.**
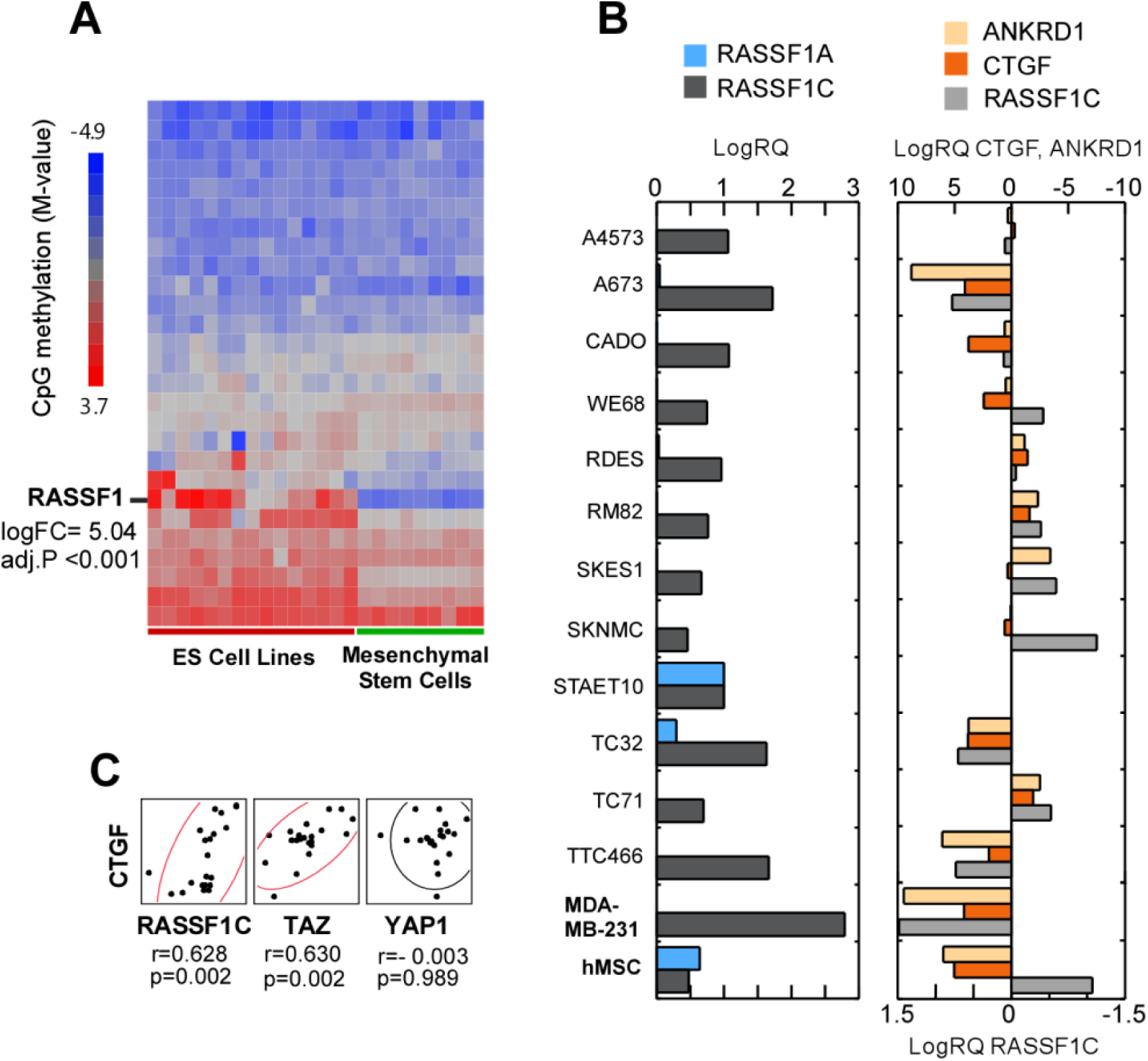
DNA methylation profiling of EwS cell lines and MSCs revealed differential CpG methylation in *RASSF1* locus. (**A**) Heat map depicting CpG methylation levels of Hippo-related loci across a panel of EwS cell lines and hMSC from EwS patients and healthy donors. (**B**) Relative quantification by qPCR of *RASS1A* and *RASSF1C* transcripts and TAZ/YAP1 target genes in a panel of EwS cell lines. A basal breast cancer cell line and hMSC are included as controls (experiments were performed with 3 biological samples in triplicates). (**C**) Correlation analyses of mRNA expression levels (qPCR) of CTGF with RASSF1C, TAZ and YAP1 (r, Pearson’s correlation coefficient).

There is extensive evidence that Src can promote YAP/TAZ activity through a variety of mechanisms, i.e. Src, and other SFKs can directly phosphorylate YAP1 and TAZ promoting their activity and stability [31]. Therefore, since *RASSF1C* activates SFKs in *RASSF1*-methylated cells, we blocked SFK activity by exposing EwS cells to dasatinib. Inhibition of SFKs resulted in reduced cell viability (SK-N-MC IC50=6,55 μM; TTC-466 IC50=2,11 μM) and upregulation of YAP1/TAZ target genes (**Fig. 5A**). Upon dasatinib treatment mRNA levels of YAP1 and TAZ remained unaffected, but TAZ protein expression was decreased and YAP1 inactivating phosphorylation increased in both cell lines (**Fig. 5A**). As an alternative approach of pharmacologic blockade of YAP1/TAZ activity we tested pitavastatin. Statins prevent nuclear localization of YAP1/TAZ via inhibition of the enzyme HMG-CoA reductase, ultimately affecting the metabolic control of YAP1/TAZ by the mevalonate pathway [32]. We also observed antiproliferative effect upon pitavastatin treatment (SK-N-MC IC50= 1,83 μM; TTC-466 IC50=1,86 μM), with mild reduction of YAP1/TAZ target genes and TAZ protein downregulation (**Fig 5A**). Neither dasatinib nor pitavastatin treatments affected EWS-FLI1 expression in SK-N-MC cell line, thus precluding the antiproliferative effect of these drugs to be mediated by the fusion protein.

**Fig 5.**
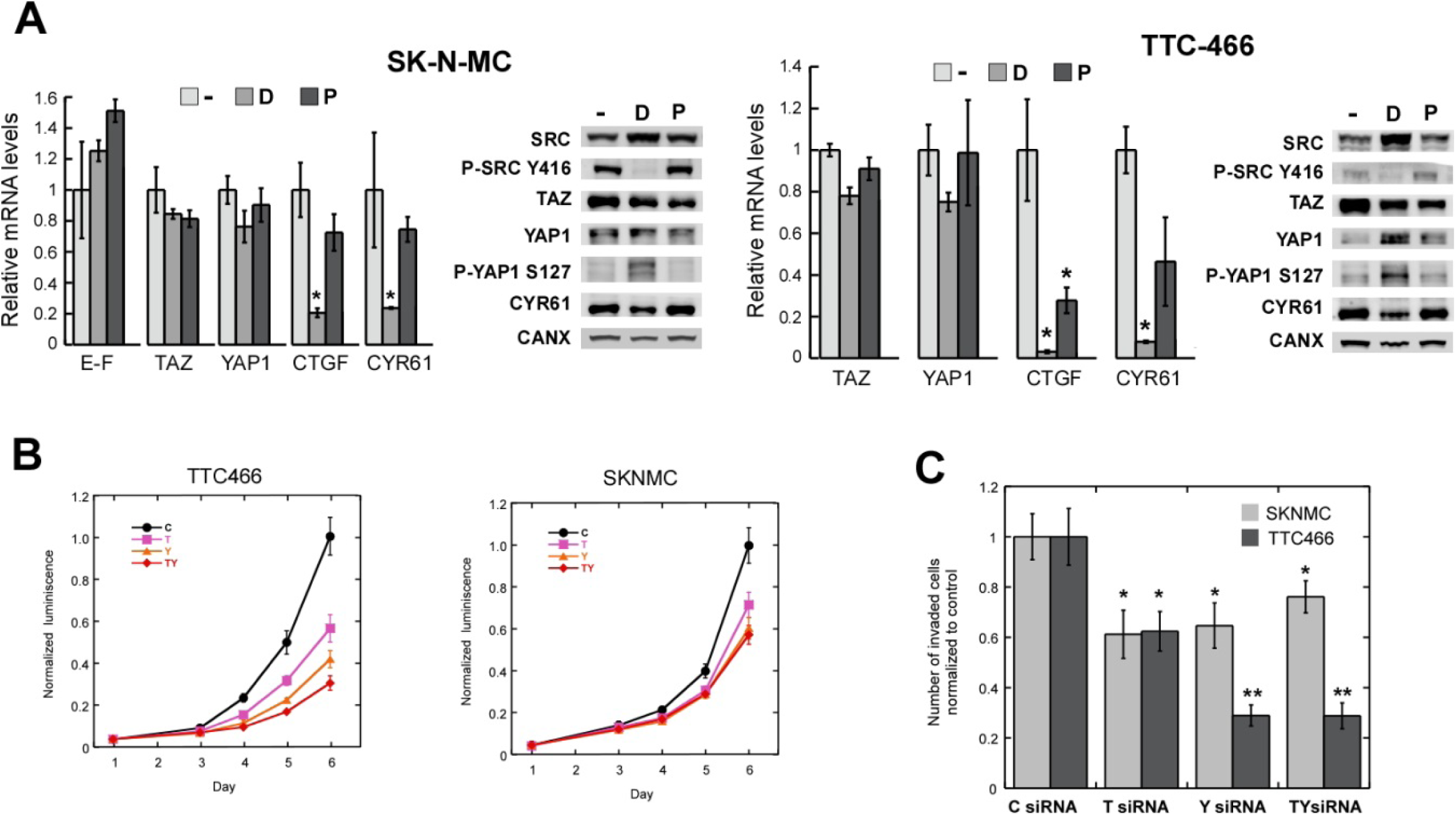
Pharmacologic inhibition and siRNA silencing of YAP1/TAZ in EwS cells. (**A**) SK-N-MC and TTC-466 cell lines were treated with Dasatinib (D, 1 uM) or Pitavastatin (P, 1 uM) during 24h, and mRNA levels of YAP1, TAZ and their target genes CTGF and CYR61 were quantified by qPCR. mRNA levels of EWS-FLI1 were evaluated in SK-N-MC cell line. Whole cell extracts were also analyzed by WB (experiments were performed with 3 biological samples in tripli cates; *p<0.05). (**B**) Proliferation curves of EwS cell lines transfected with control siRNA (C), siRNA targeting YAP1 (siY), TAZ (siT) or a combination of siRNAs to deplete both factors simultaneously (siYT). Two different siRNAs were used to knockdown each factor rendering similar levels of silencing (**Suppl. Fig. 4**), data are only shown for one of the siRNAs. Results are expressed as the mean ± SD of three independent experiments performed in triplicate. All the conditions were significantly different form the control (p<0.05) (**C**) Invasion assay of EwS cell lines upon individual or combined silencing of YAP1 and TAZ (*p<0.05, **p<0.005).

### YAP1/TAZ loss-of-function affects cell proliferation and invasion capacity in EwS cells

To assess the oncogenic properties of YAP1 and TAZ in EwS cells, we induced transient knockdown of YAP1, TAZ or simultaneous depletion of both factors, and evaluated cell proliferation, invasion and migration capacity of the silenced cells. We observed inhibition of proliferation in knockdown cells for every individual or combined siRNA transfection. Individual depletion of YAP1 inhibited cell growth more efficiently than TAZ silencing (**Fig 5B**). YAP1/TAZ silenced cells showed a significantly reduced invasive capacity as well (**Fig 5C**). Migration capacity of EwS cells upon YAP1/TAZ silencing was not significantly altered as compared to the control, but a slight trend towards diminished migration was observed in the doble-silenced cells (**Suppl. Fig. 6**).

### YAP1/TAZ transcription activity is anti-correlated with EWS-FLI1 transcriptional signature

To evaluate the transcriptome modulation by YAP1/TAZ we conducted gene expression profiling by Affymetrix microarrays in SK-N-MC cells upon simultaneous silencing of both factors. We observed differential expression of 938 coding genes (**Suppl. Table 3**) including well-stablished YAP1/TAZ target genes, such as *CYR61*, *CTGF* or *AMOT*, which were confirmed by qPCR analyses in two EwS cell lines with different gene fusions (**Fig. 6A**). Similar results were obtained with individual silencing of each factor (**Suppl. Fig. 5**). Of note, expression levels of EWS-FLI1 were not affected in SK-N-MC (**Fig. 6A**) and other EwS cell lines tested (**Suppl. Fig. 5**).

**Figure 6.**
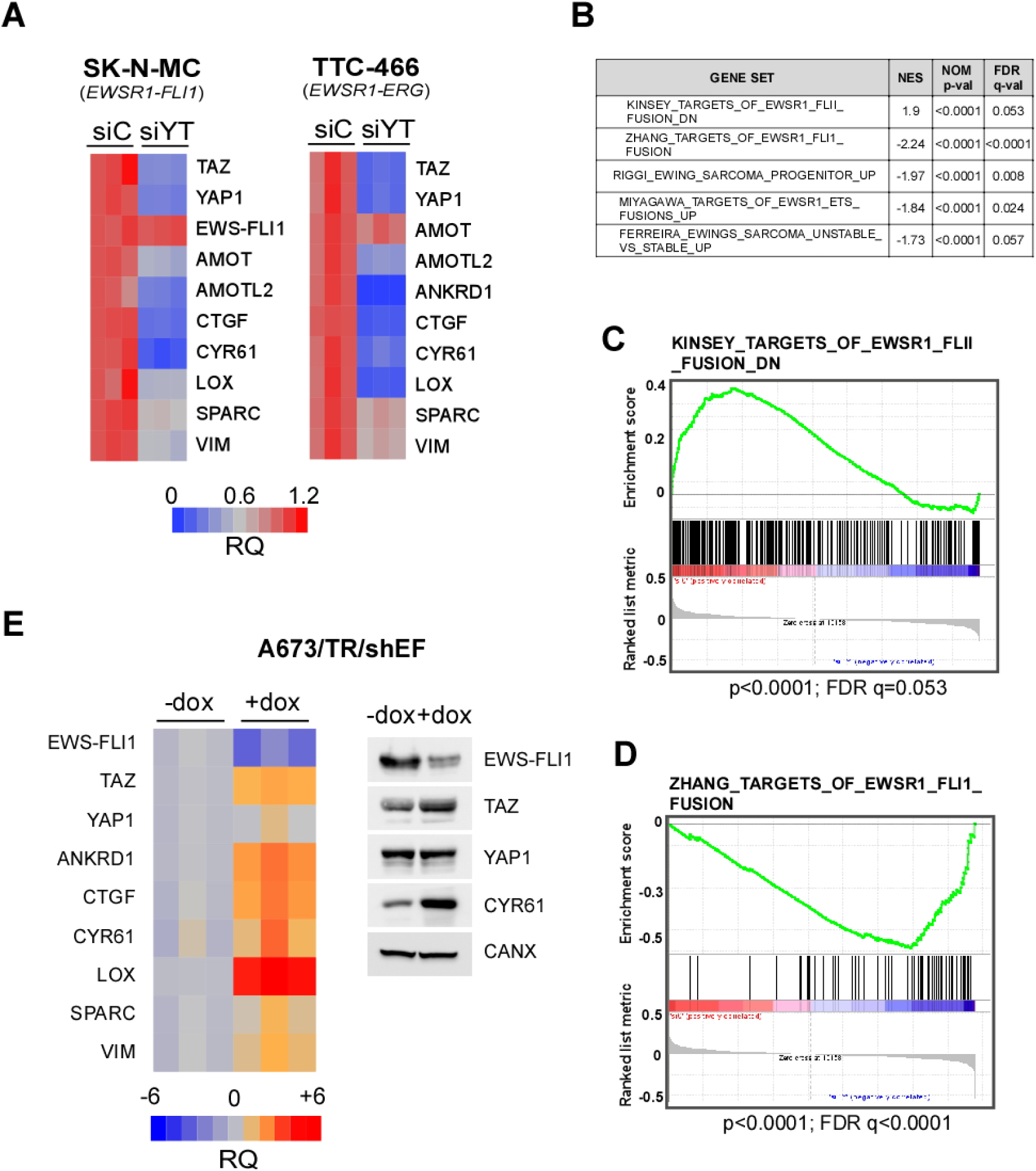
YAP1/TAZ induce an EWS-FLI1-oppossite gene signature. (**A**) qPCR assays for YAP1/TAZ target genes in EwS cell lines with different gene fusions upon siRNA depletion of YAP1 and TAZ (see Suppl Fig. x. for qPCR with individual silencing of each factor). (**B**) EwS gene sets with a positive and negative enrichment score for YAP1/TAZ regulated genes in SK-N-MC cell line. (**C**, **D**) Examples of YAP1/TAZ rank-ordered target genes compared with downregulated and upregulated EWS-FLI1 gene sets respectively (NES, normalized enrichment score). (**E**) qPCR and WB assays showing derepression of TAZ and YAP1/TAZ target genes upon silencing of EWS-FLI1 in the cell line A673 (dox, doxycycline induction of shRNA targeting EWS-FLI1).

Next, we collated this transcriptional profile with previously published curated gene sets. Interestingly, we found significant enrichment for several EwS-related gene signatures both in YAP1/TAZ-correlated and anticorrelated genes (**Fig. 6C**). YAP1/TAZ-anticorrelated genes were significantly over-represented among EwS induced gene sets, and inversely YAP1/TAZ-correlated genes overlapped with EwS repressed genes. Thus, suggesting opposite transcriptional activity of EWS-FLI1 fusion gene and YAP1/TAZ factors. Accordingly, depletion of the EWS-FLI1 protein in the A673 EwS cell line resulted in the induction of YAP1/TAZ-regulated genes, as well as TAZ but not YAP1 factor (**Fig. 6C, D**). Therefore, transcriptional antagonism may be partially explained by EWS-FLI1-mediated downregulation of TAZ. We confirmed these observations in public datasets for EWS-FLI1 silencing in five EwS cell lines [33], and for ectopic expression of EWS-FLI1 in embryonic stem cells [34] (**Suppl. Fig. 7)**. These observations are in accordance with recent reports describing that several genes are inversely regulated by TEAD factors and EWS-FLI1 [35, 36]. TEADs are the main transcription factors partners of YAP1 and TAZ, and usually associate with AP-1 transcription factors at distal enhancers [28, 37]. Both TEAD and AP-1 conserved binding motifs are present in EWS-FLI1 regulated genes [35]. Furthermore, EWS-FLI1 binding at *WWTR1* locus coding for TAZ correlates with a decrease of TAZ mRNA expression, suggesting direct repression of TAZ by EWS-FLI1 (**Suppl. Fig. 8**).

## Discussion

In the present study, we have shown that YAP1/TAZ expression associates with disease progression and poor prognosis in a large retrospective series of EwS patients. Few reports have addressed this issue so far, and the reported series were smaller, i.e. Ahmed AA. et al. [38] observed that YAP1 expression can be detected in 47% of samples (in a series of 32 cases) without association with survival, whereas in another study with only 5 cases, 60% and 80% showed YAP1 and TAZ expression respectively [39]. Other pediatric sarcomas such us rhabdomyosarcoma, osteosarcoma or neuroblastoma have been reported to express YAP1 and TAZ, with an impact in patient prognosis and conferring resistance to current therapies [40–44]. Another fact that supports the relevance of YAP1/TAZ and other Hippo signaling effectors in sarcomas is their involvement in recurrent fusion genes in certain histological types, such as epithelioid hemangioendothelioma (*WWTR1-CAMTA1*, *YAP1-TFE3*), epithelioid haemangioma (*WWTR1-FOSB*) or spindle cell rhabdomyosarcoma (*VGLL2–CITED2*, *VGLL2–NCOA2*, *TEAD1–NCOA2*)[45, 46]. Notwithstanding, aberrant activation of YAP1/TAZ in cancer is often promoted by mechanisms not involving somatic alterations. We have observed that epigenetic regulation of the *RASSF1* locus could affect the expression of YAP1/TAZ target genes in EwS cell lines (**Fig. 4**). This result may explain previous observations describing a correlation of hypermethylation of *RASSF1* and *RASSF2* with worse clinical outcome in EwS [47, 48]. Moreover, Src kinase activation of invadopodia in response to stress in EwS [49] could be related to SFK-mediated activation of YAP1/TAZ by *RASSF1C* (**Fig. 5A**). However, YAP1/TAZ activation does not seem to rely on *RASSF1* hypermethylation in hMSC (**Fig. 4**), the putative cell of origin of EwS, which exhibits high expression levels of YAP1 and TAZ (**Fig. 1**). Unaffected expression levels of YAP1 and derepression of TAZ upon EWS-FLI1 silencing (**Fig. 6C, D**) also supports the notion that both factors are maybe expressed in the cell of origin, as proposed for ZEB2, an EMT (epithelial–mesenchymal transition) inducer like YAP1 and TAZ [50].

The association of YAP1/TAZ with metastatic spread could be arguably related to the relative levels of the fusion protein, recently reported to promote phenotypic plasticity of EwS cells [16]. In this scenario, YAP1/TAZ may promote a mesenchymal phenotype in EWS-FLI1 depleted EwS cells together with Wnt/beta-catenin [51], since it is well-established that the crosstalk between Hippo and wnt signaling is essential for tumor progression in several types of cancer [52]. As it has been described for Wnt/beta-catenin [51], the opposing transcriptional signature between YAP1/TAZ and EWS-FLI1 could partly contribute to the metastatic process. I.e. We found strong downregulation or upregulation of LOX (a mediator of metastasis [16]) in YAP1/TAZ-silenced or EWS-FLI1-silenced cells respectively. These results suggest that LOX expression in EwS could result both of derepression in a low-level state of the fusion protein as well as of inducer mechanisms involving YAP1 or TAZ. In line with this, ChIP-seq data from Bilke S. et al. [53] reveal that EWSR1-FLI1 binds at regulatory elements of some of the well-established TAZ/YAP1 target genes [36]. Furthermore, anticorrelation of AP-1 induced genes and EWSR1-FLI1 transcriptional signature was observed in the same cell model that we used in this work: inducible silencing of EWSR1-FLI1 in A673 cell line [36]. It is well established that YAP1/TAZ/TEAD transcriptional complexes usually cooperate with AP-1 at regulatory DNA modules to synergistically activate target genes [28, 37]. Therefore, the transcriptional antagonism might be consequence of some interference between YAP1/TAZ/TEAD-AP1 complexes and the fusion protein, as demonstrated by Katschnig et al. [35]. Another mechanism contributing to the opposing gene signatures might involve Ewing sarcoma-associated transcript 1 (EWSAT1), which was found to be significantly induced in YAP1/TAZ-silenced SK-N-MC cells (**Suppl. Table 3**). EWSAT1 is a long noncoding RNA that mediates EWS-FLI1 gene repression via interaction with a heterogeneous nuclear ribonucleoprotein [54]. In addition, we have observed inhibition of TAZ expression associated to the presence of EWS-FLI1, which also binds DNA at *WWTR1* locus (**Suppl. Fig. 8**). Indeed, regulation of TAZ seems to occur at the transcriptional level, whereas YAP1 activity is not correlated with mRNA levels (**Figs. 4C, 6E**).

In summary, our study reveals that the interplay between Hippo pathway effectors YAP1/TAZ and the function of the gene fusion is relevant to shape the transcriptional program in EwS. The transcriptional output elicited by these factors deserves further characterization since our observations provide clinical evidence that YAP1/TAZ expression associates with disease progression in EwS patients. Studies with larger prospective series are needed in order to corroborate our observations and to stablish whether YAP1/TAZ could serve as reliable biomarkers to stratify and identify patients who could benefit from targeted therapies.

## Acknowledgments

This research has been conducted using samples from the Hospital Universitario Virgen del Rocío-Instituto de Biomedicina de Sevilla Biobank (Andalusian Public Health System Biobank and ISCIII-Red de Biobancos PT17/0015/0041). The authors thank the donors for the human specimens used in this study. This work was supported by a grant from the Fundación Pública Andaluza Progreso y Salud (Junta de Andalucía) and JANSSEN CILAG, S.A. (Grant No. PI-0344-2014) to JDM. PRN is a PhD student recipient of a PFIS fellowship to Enrique de Alava (Grant No. F109/00193). JDM, LRP and ATM are PhD researchers funded by the Asociación Española Contra el Cáncer (aecc, GCB13-1578). CJ works as a laboratory technician supported by the ISCIII. EDA’s lab is supported by the AECC project (GCB13-1578); ISCIII-FEDER (PI14/01466); CIBERONC (CB16/12/00361), Asociación Pablo Ugarte and Fundación María García Estrada. The laboratory of TGPG is supported supported by the ‘Verein zur Förderung von Wissenschaft und Forschung an der Medizinischen Fakultät der LMU München’ (WiFoMed), by LMU Munich’s Institutional Strategy LMUexcellent within the framework of the German Excellence Initiative, the ‘Mehr LEBEN für krebskranke Kinder – Bettina-Bräu-Stiftung’ (to TGPG), the Fritz-Thyssen Foundation (FTF-40.15.0.030MN), the Kind-Philipp-Foundation, the Matthias-Lackas Foundation, the Dr. Leopold and Carmen Ellinger Foundation, the Wilhelm-Sander-Foundation (2016.167.1), the German Cancer Aid (DKH-70112257), the Gert und Susanna Mayer Foundation, and the Deutsche Forschungsgemeinschaft (DFG-391665916).

The authors thank Dr. Javier Alonso for providing the cell line A673 transfected with a inducible shRNA against EWS-FLI1. We also thank Dr. Stefano Piccolo for the luciferase reporter plasmid with TEAD motifs (Addgene plasmid # 34615), Dr. Mark Bond for providing the plasmid pTNT-min, and D. Campana for hMSC TERT cell line.

## Author contributions

JDM, PRN and LRP contribute equally to this work and were responsible for the experimental design and the undertaking of the experiments. EA and DM reviewed the pathologic and immunohistochemical analyses, and the clinical data. ATM, PPC and CJ carried out some of the experiments. JDM, LRP, and PRN performed the statistical analysis and interpreted the data. TGPG analyzed public datasets. JDM designed the study, and LRP, EA and JDM were involved in writing the paper. JDM generated the figures and drafted the manuscript. All authors contributed to the editing of the manuscript and gave their approval of the final version.

---

**Suppl. Fig. 1.**
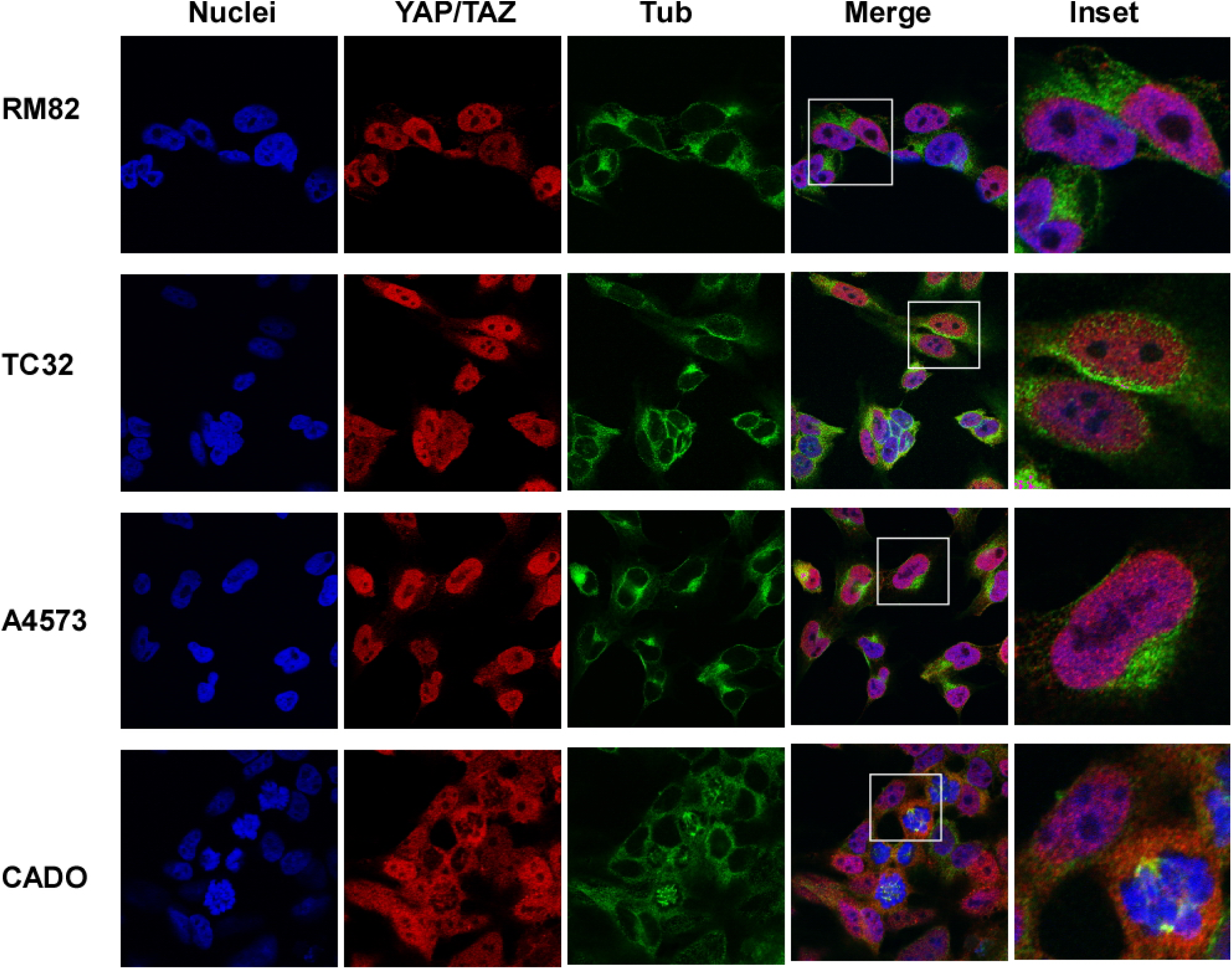
Immunofluorescence microscopy with the indicated antibodies (60X).

**Suppl. Fig. 2.**
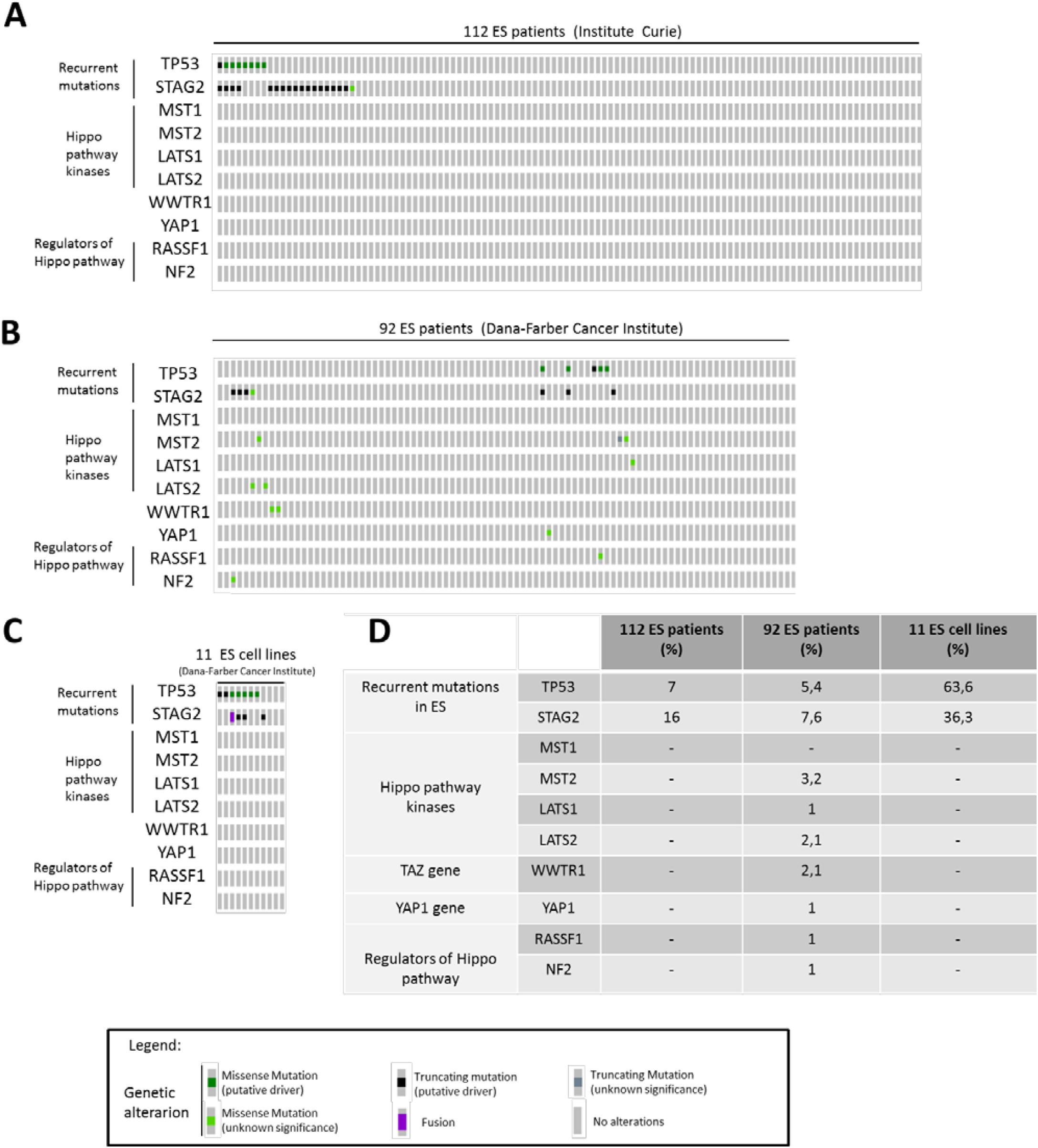
Hippo pathway-associated genes do not harbor recurrent mutations in EwS cell lines and patients. Mutational profile in different gene sets across 112 ES patients (Institut Curie; PMID: 25223734)(A) 92 EwS patients (Dana-Farber Cancer Institute; PMID: 25186949) (B) and 11 EwS cell lines (Dana-Farber Cancer Institute; PMID: 25186949) (C). Cell lines were, in order: A673, TC71, TTC466, EW8, EWS834, EWS502, RDES, TC32, CHLA258, CADO-ES and SKNEP1. Each EwS patient/cell line is represented by a grey bar. D. Genetic mutation frequency (%) at indicated gene sets across different studies.

**Suppl. Fig. 3.**
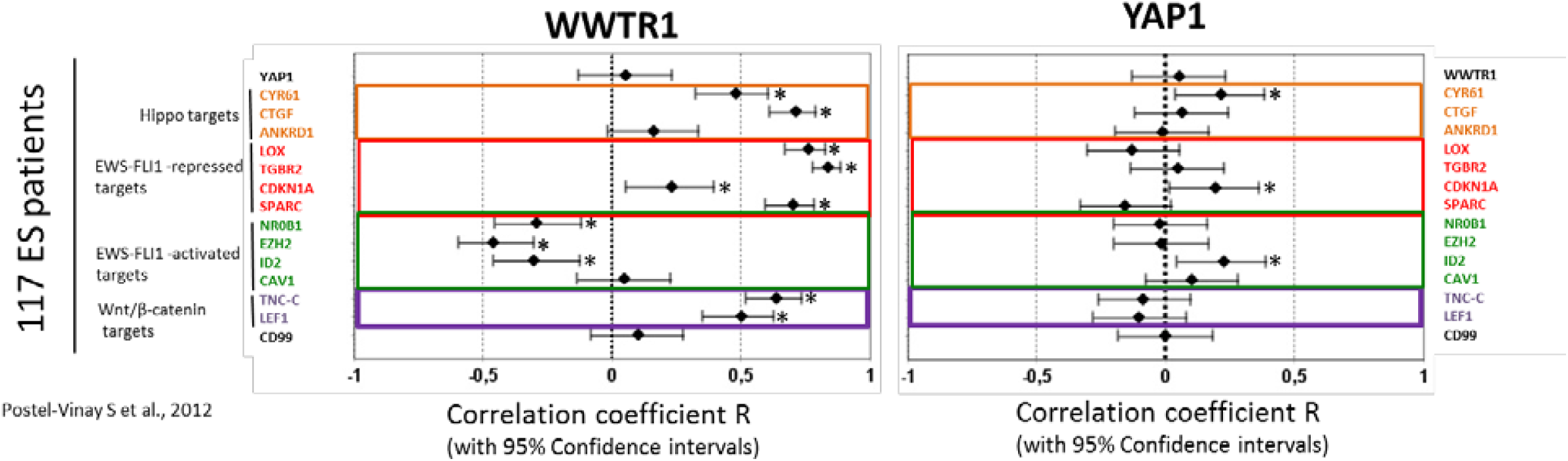
WWTR1 (TAZ) expression correlates with EWS-FLI1 targets in EwS patients.

**Suppl. Fig. 4.**
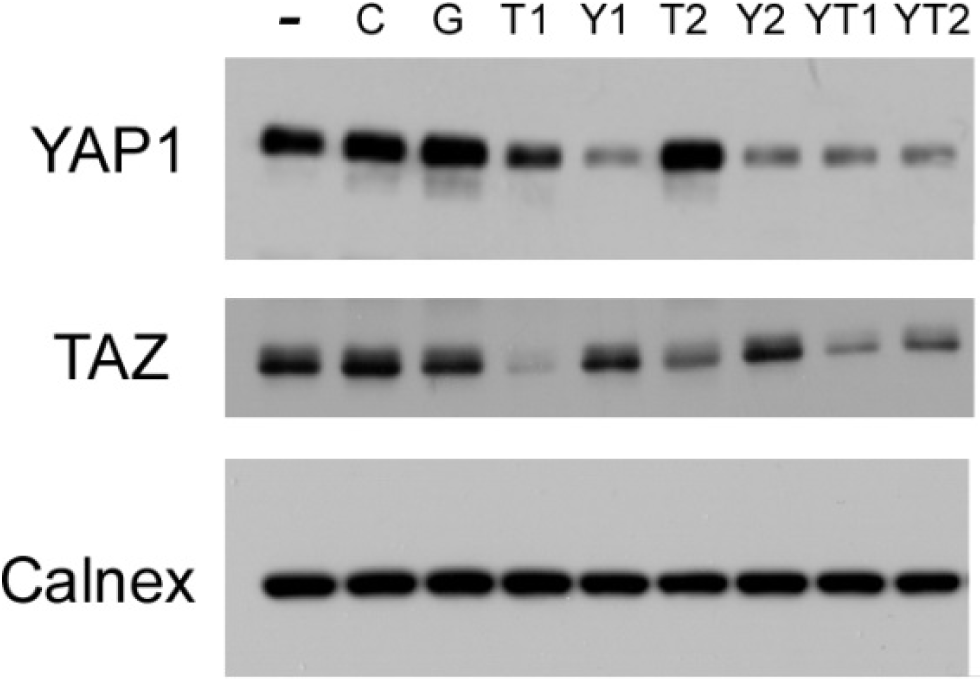
Western blot assay to test silencing of YAP1 and TAZ in SK-N-MC cell line with two different siRNAs in individual or doble trasfection. -, non transfected; C Sramble siRNA; G, siRNA targeting GAPDH; T1 and T2, siRNAs targeting TAZ; Y1 and Y2, siRNAs targeting YAP1; YT1, combination of siRNAs T1 and Y1; YT2, combination of siRNAs T2 and Y2.

**Suppl. Fig. 5.**
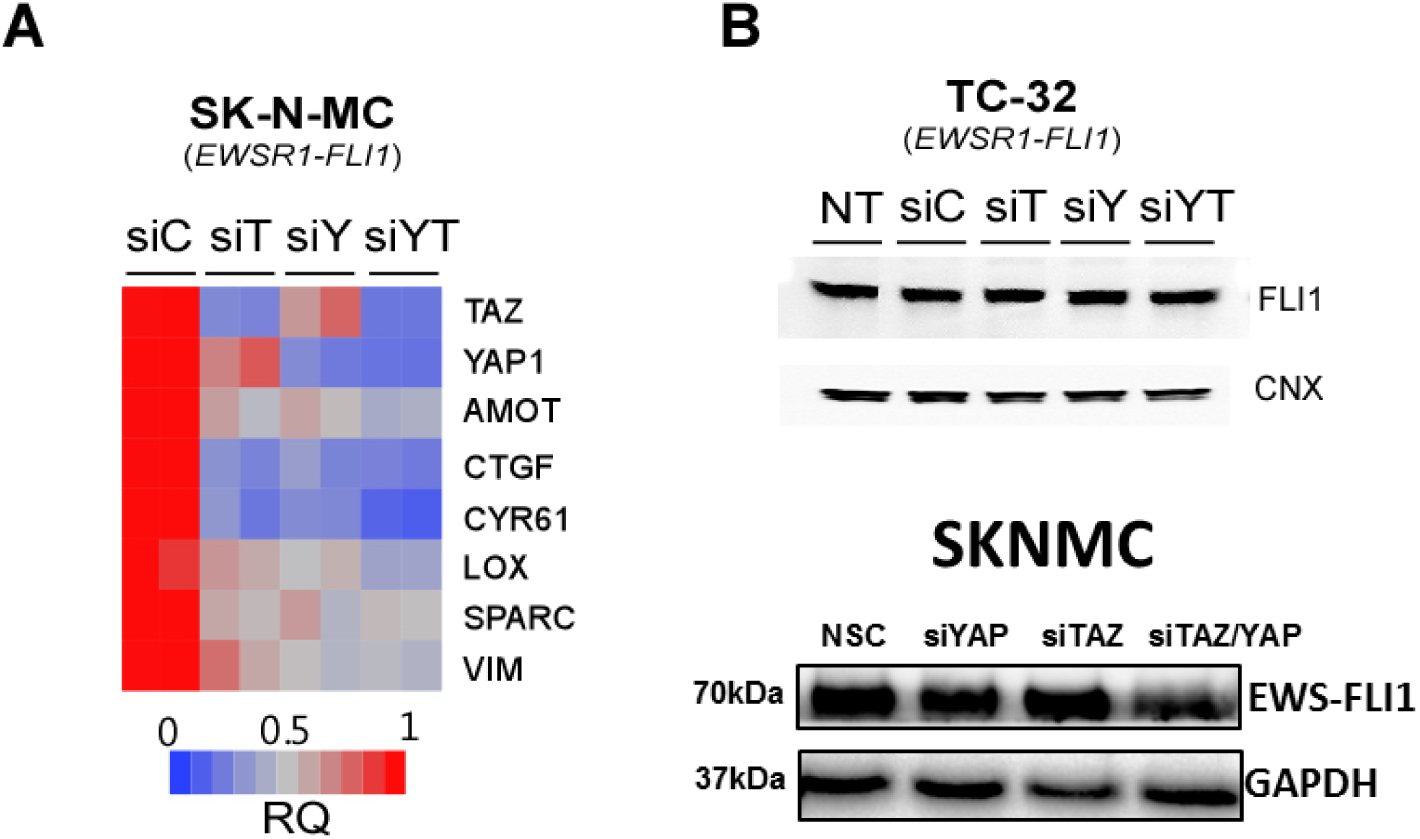
(**A**) qPCR for quantification of YAZ1/TAZ target genes in SK-N-MC cell line upon siRNA silencing of YAP1 and TAZ. (**B**) Lysates from TC-32 and SK-N-MC silenced cells were probed with anti-FLI1 to evaluate EWS-FLI1 protein expression.

**Suppl. Fig. 6.**
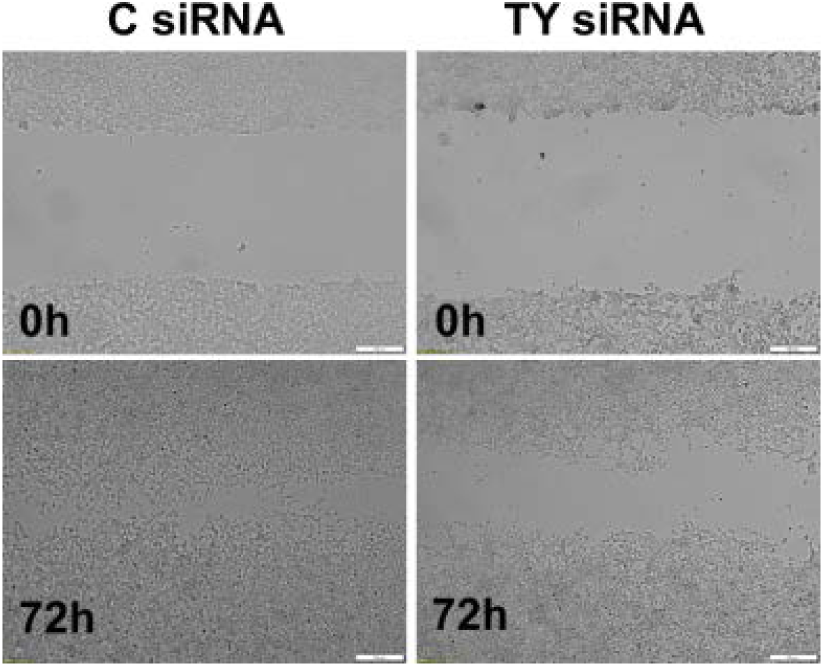
Representative images of a wound healing assay using control or YAP1/TAZ-silenced TTC-466 cells.

**Supp Fig. 7.**
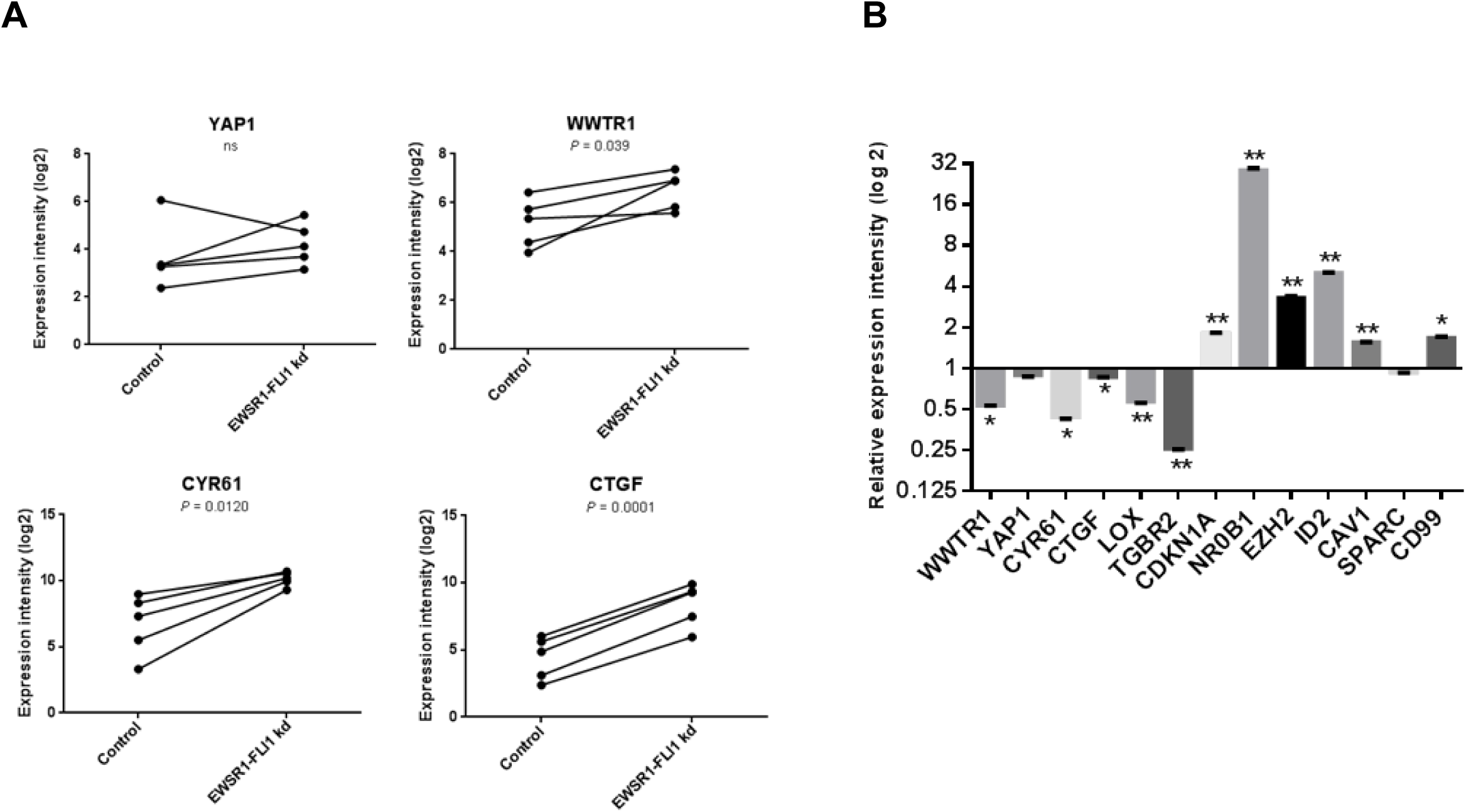
EWS-FLI1 modulates WWTR1 (TAZ) and Hippo target genes expression in different in vitro models. **A.** Analysis of gene expression levels of YAP1, WWTR1, CYR61 and CTGF after shRNA-mediated knockdown (kd) of EWSR1-FLI1 in 5 ES cell lines (WE68, SK-N-MC, TC252, STA-ET-1, STA-ET-7.2) (GSE14543). Data are represented in a before-after plot in which each dot represents a cell line. Two-tailed student’s t test. **B.** Analysis of gene expression levels of WWTR1, YAP1, TAZ/YAP1 targets (CYR61, CTGF), EWS-FLI1-repressed (LOX, TGFBRII, CDKN1A) and - induced targets (NR0B1, EZH2, ID2, CAV1, SPARC) and CD99 expression in human embryonic stem cells after ectopic expression of EWSR1-FLI1 (GSE64686). Mean±S.E.M are depicted of the three biological triplicates for each gene. Two-tailed student’s t test.

**Suppl. Fig. 8.**
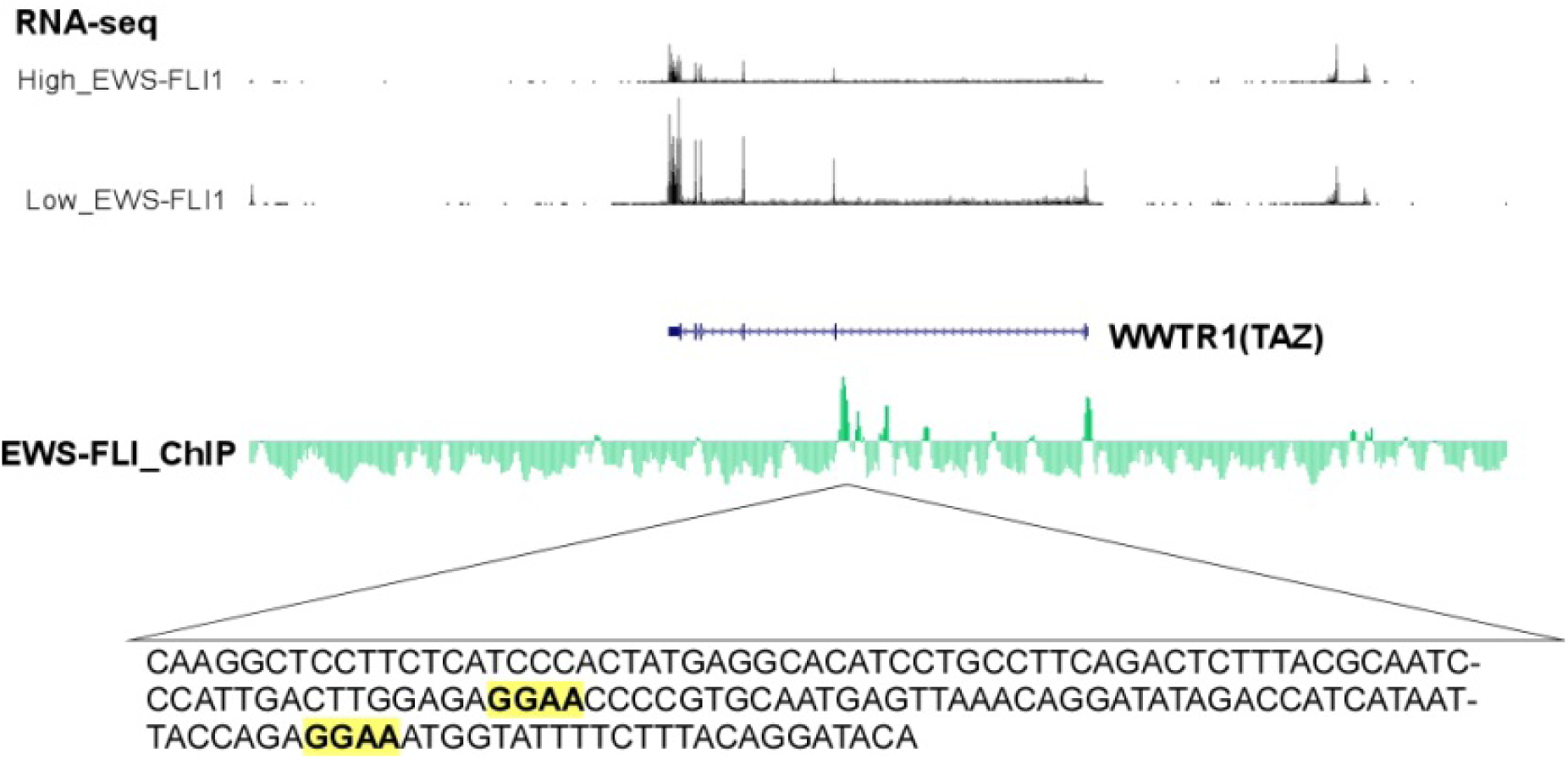
Genome browser screenshot of *WWTR1* locus showing RNA expression upon inducible silencing of EWS-FLI in A673 cell line. EWS-FLI binding peaks within the locus are depicted and nucleotide sequence is indicated for one peak with ETS binding motifs. Data were retrieved from http://tomazou2015.computational-epigenetics.org [36].

**Supplementary Table 1.**
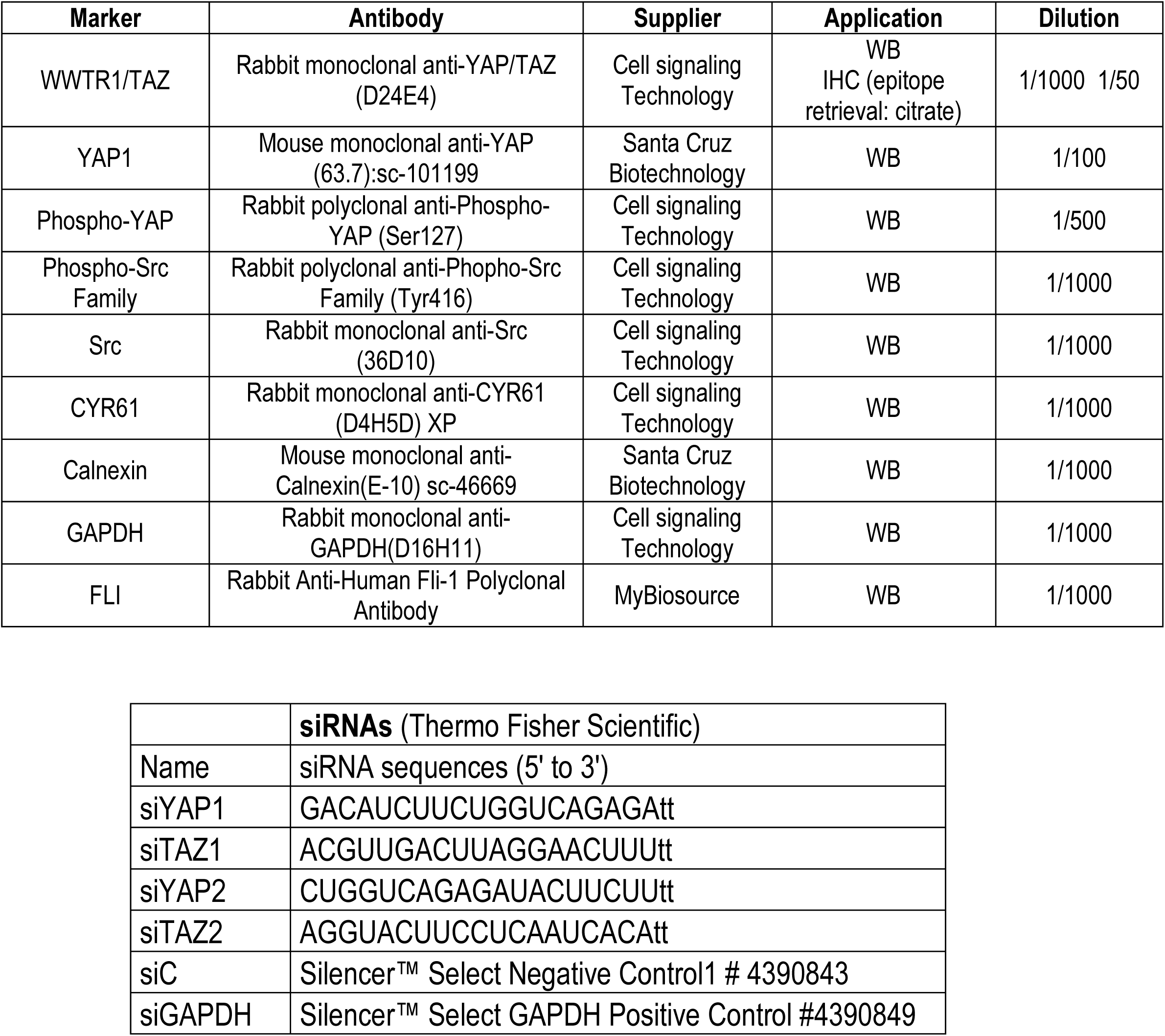
Antibodies and siRNAs, suppliers and dilutions.

**Suppl Table 2.**
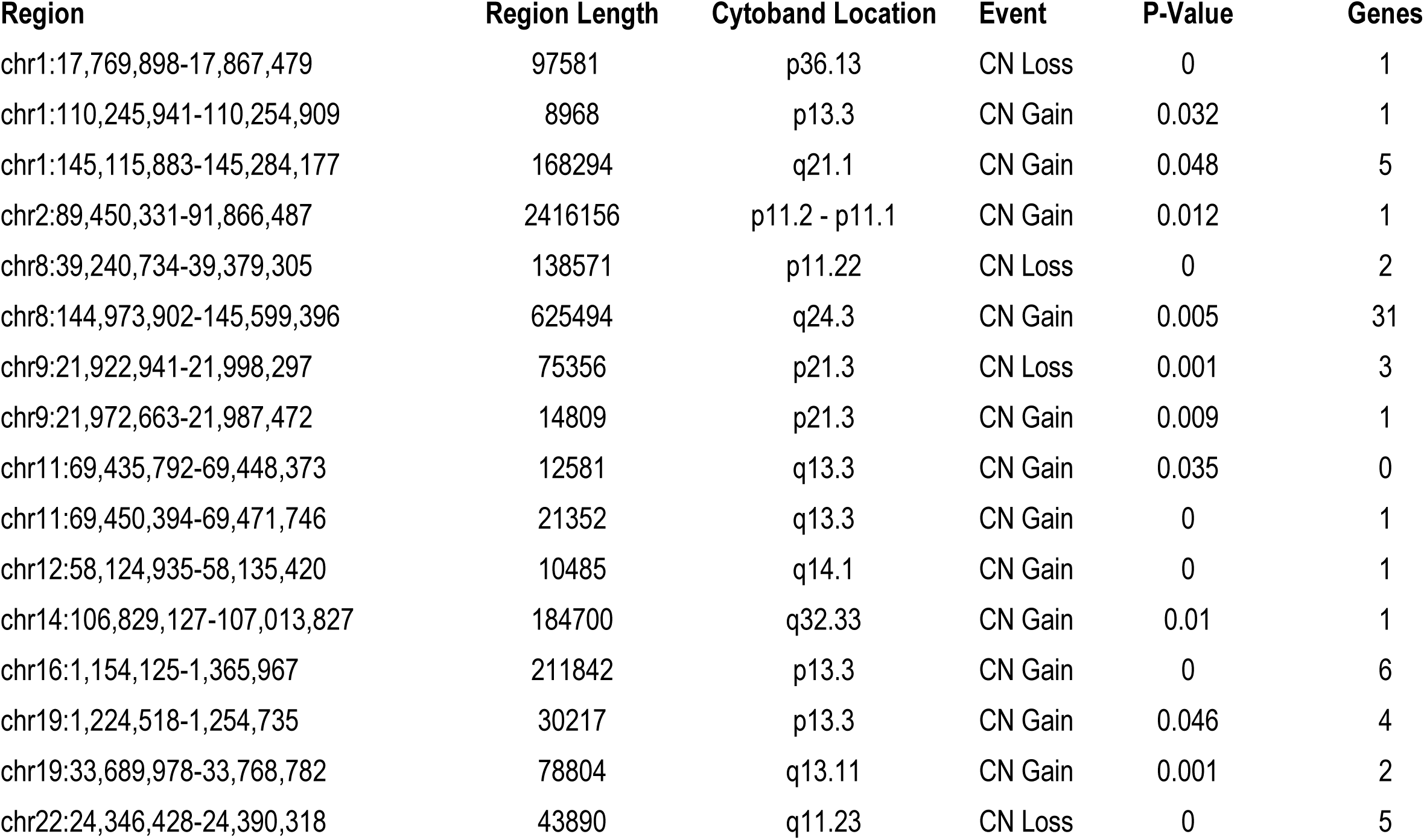
STAC peaks identified in 24 EwS samples. Displaying only significant regions with p-value less than or equal to 0.05

| Region | Region Length | Cytoband Location | Event | P-Value | Genes |
| --- | --- | --- | --- | --- | --- |
| chr1:17,769,898-17,867,479 | 97581 | p36.13 | CN Loss | 0 | 1 |
| chr1:110,245,941-110,254,909 | 8968 | p13.3 | CN Gain | 0.032 | 1 |
| chr1:145,115,883-145,284,177 | 168294 | q21.1 | CN Gain | 0.048 | 5 |
| chr2:89,450,331-91,866,487 | 2416156 | p11.2 - p11.1 | CN Gain | 0.012 | 1 |
| chr8:39,240,734-39,379,305 | 138571 | p11.22 | CN Loss | 0 | 2 |
| chr8:144,973,902-145,599,396 | 625494 | q24.3 | CN Gain | 0.005 | 31 |
| chr9:21,922,941-21,998,297 | 75356 | p21.3 | CN Loss | 0.001 | 3 |
| chr9:21,972,663-21,987,472 | 14809 | p21.3 | CN Gain | 0.009 | 1 |
| chr11:69,435,792-69,448,373 | 12581 | q13.3 | CN Gain | 0.035 | 0 |
| chr11:69,450,394-69,471,746 | 21352 | q13.3 | CN Gain | 0 | 1 |
| chr12:58,124,935-58,135,420 | 10485 | q14.1 | CN Gain | 0 | 1 |
| chr14:106,829,127-107,013,827 | 184700 | q32.33 | CN Gain | 0.01 | 1 |
| chr16:1,154,125-1,365,967 | 211842 | p13.3 | CN Gain | 0 | 6 |
| chr19:1,224,518-1,254,735 | 30217 | p13.3 | CN Gain | 0.046 | 4 |
| chr19:33,689,978-33,768,782 | 78804 | q13.11 | CN Gain | 0.001 | 2 |
| chr22:24,346,428-24,390,318 | 43890 | q11.23 | CN Loss | 0 | 5 |

**Supplementary Table 3.**
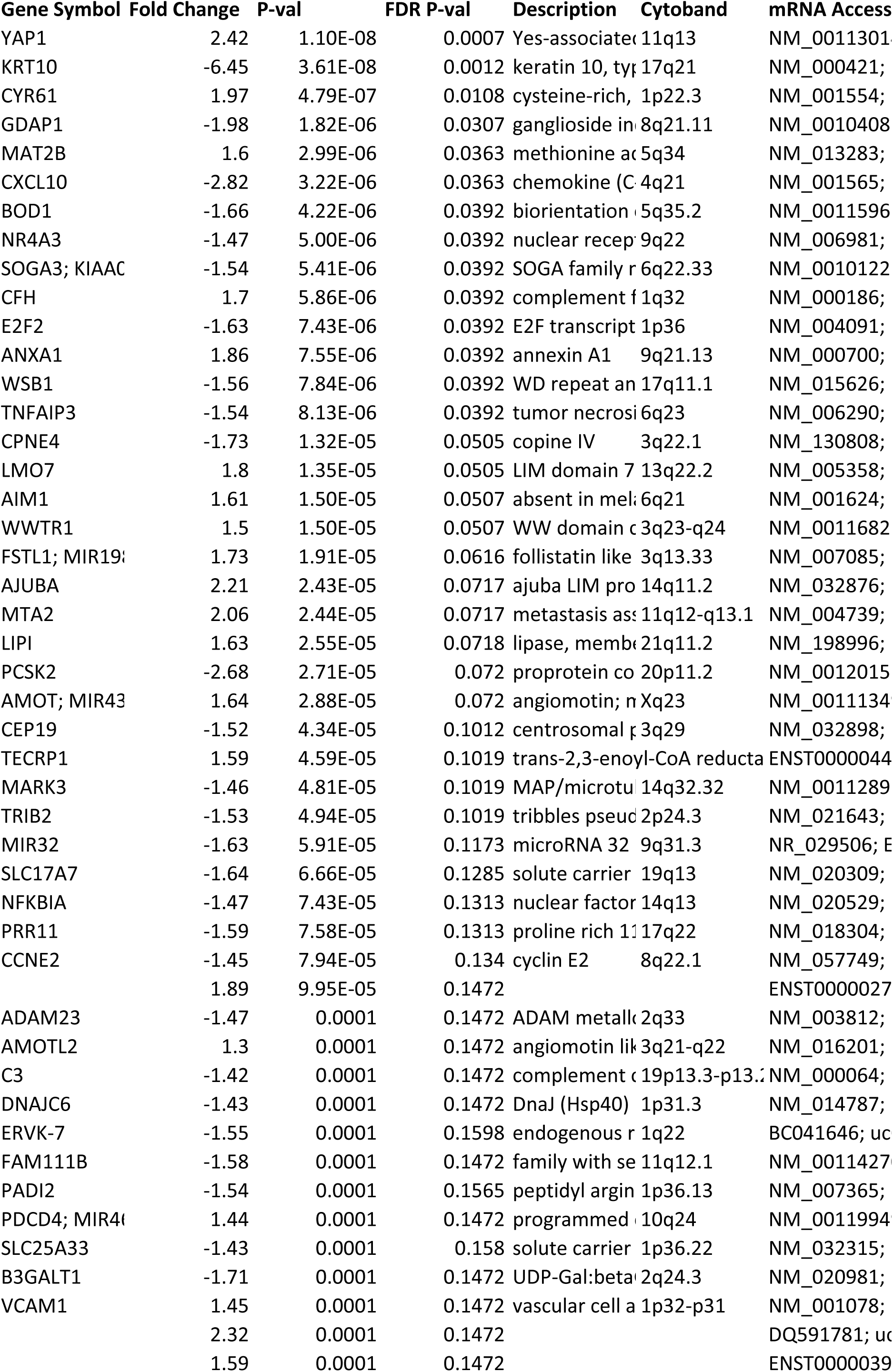
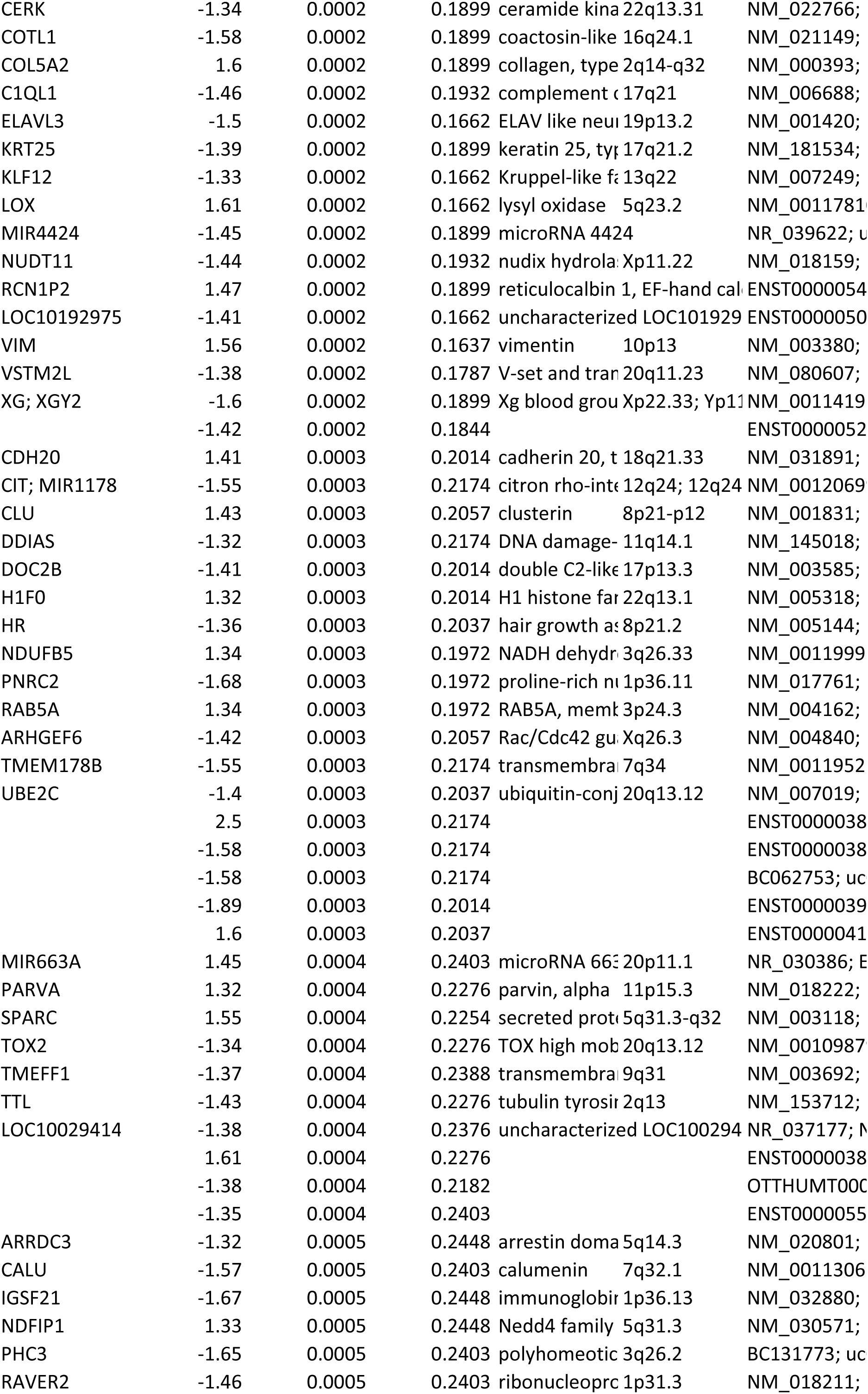
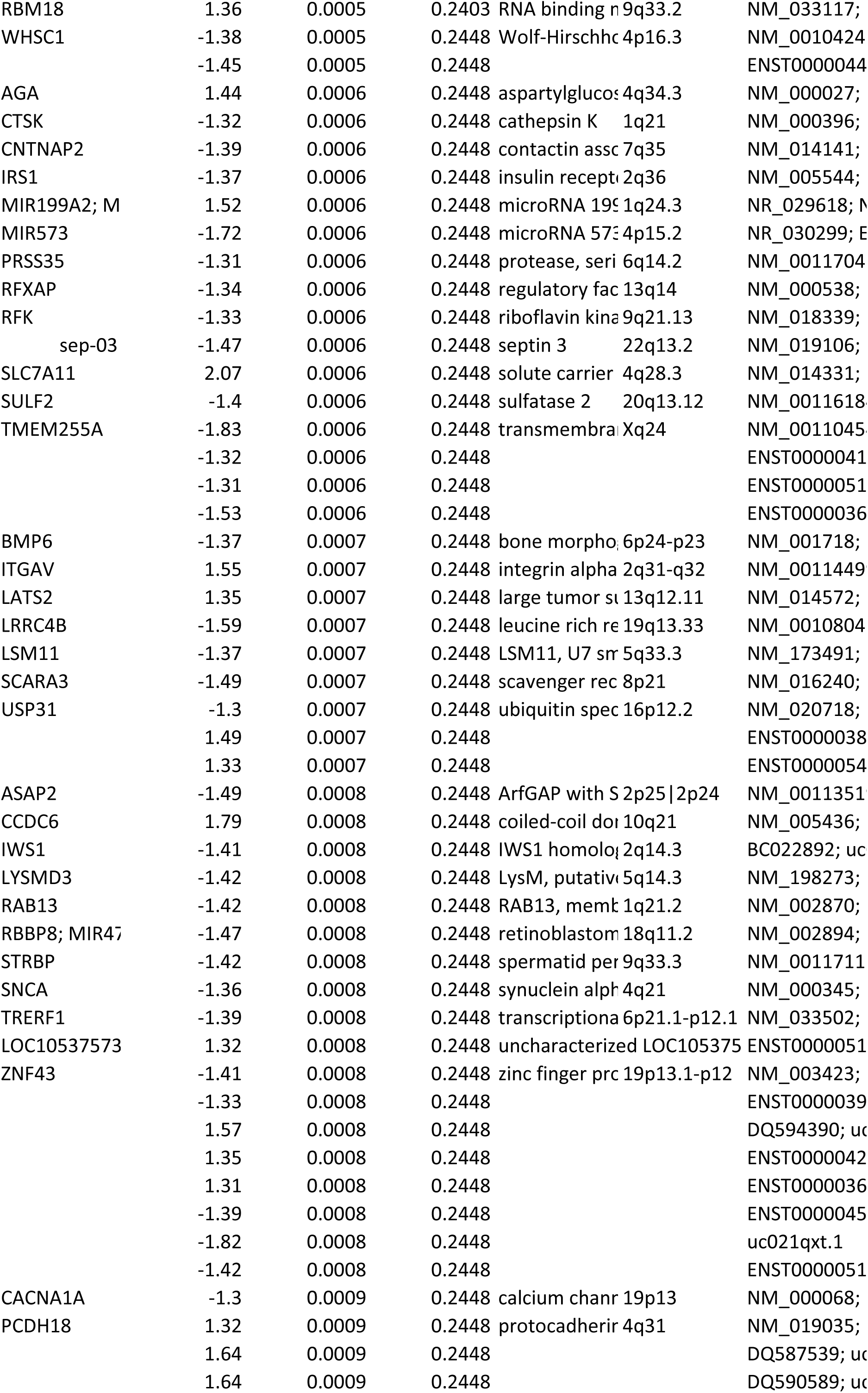
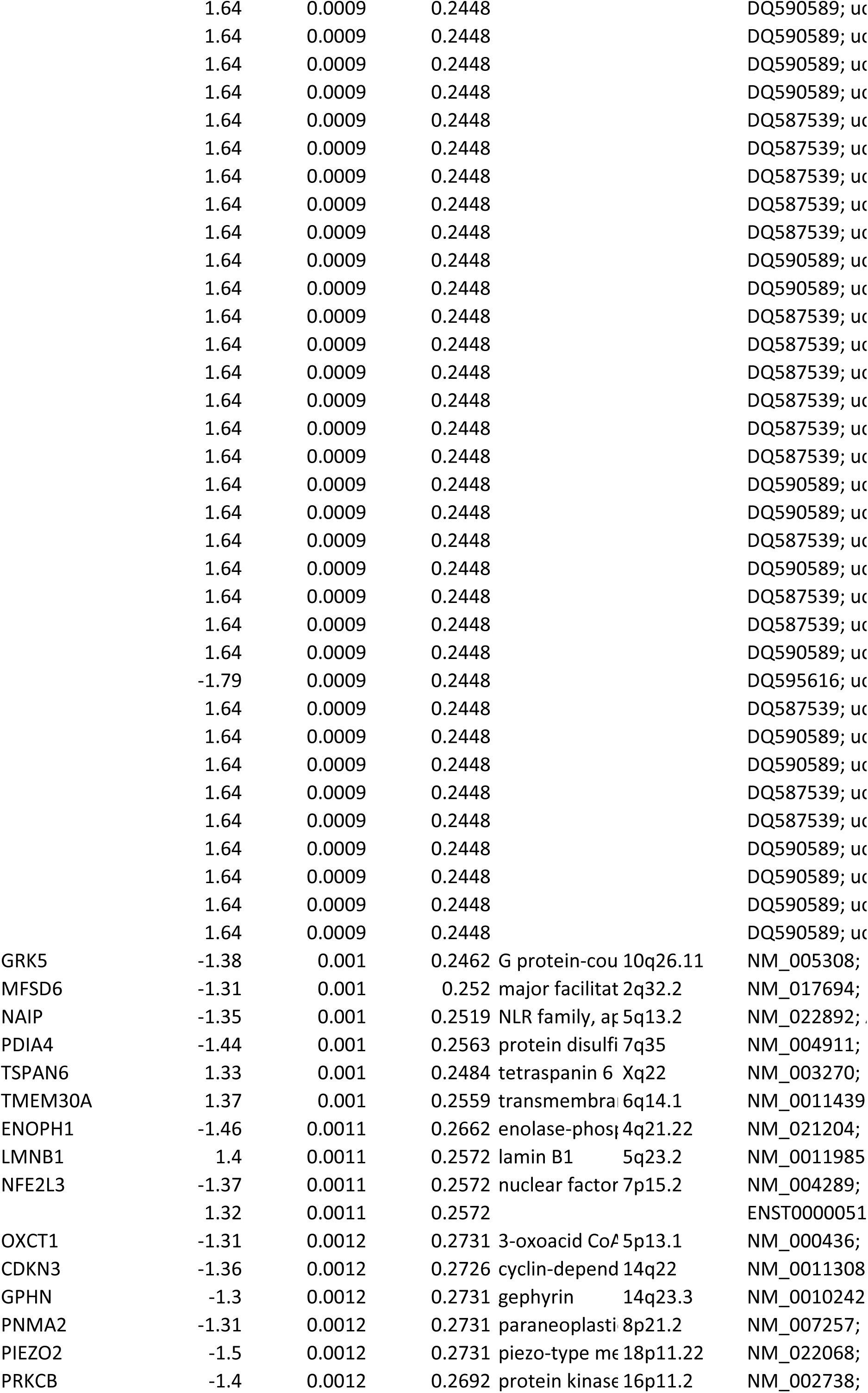
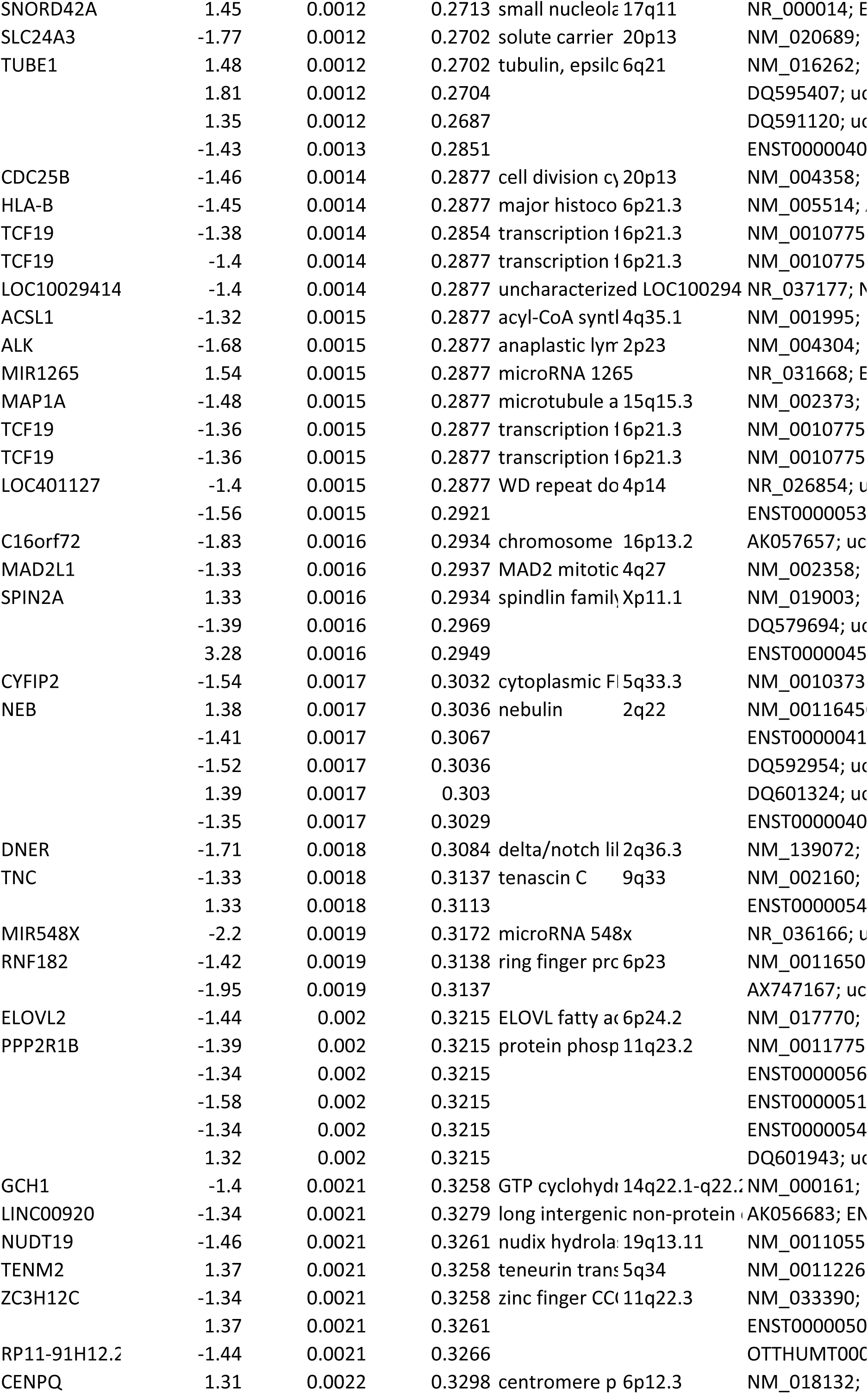
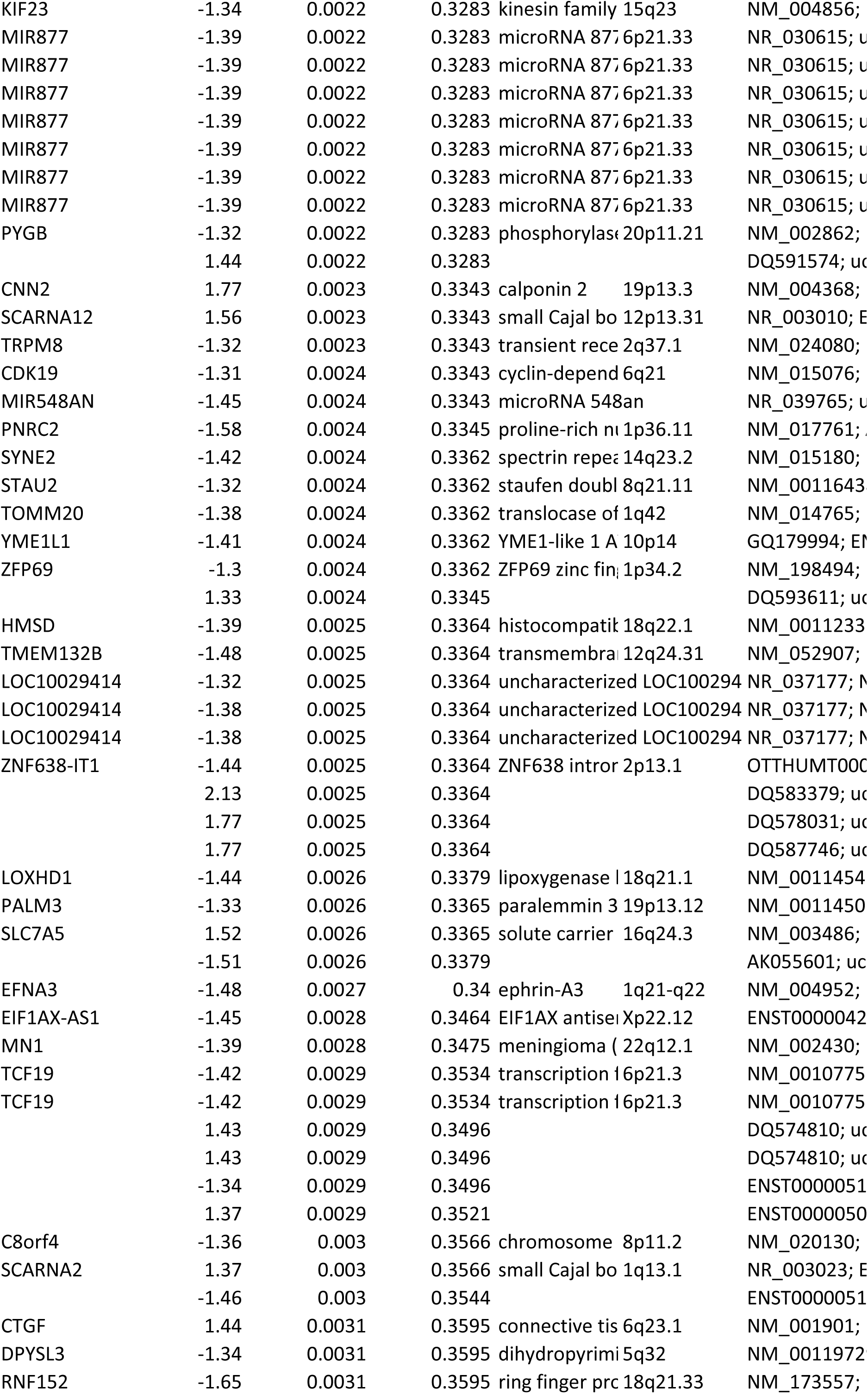
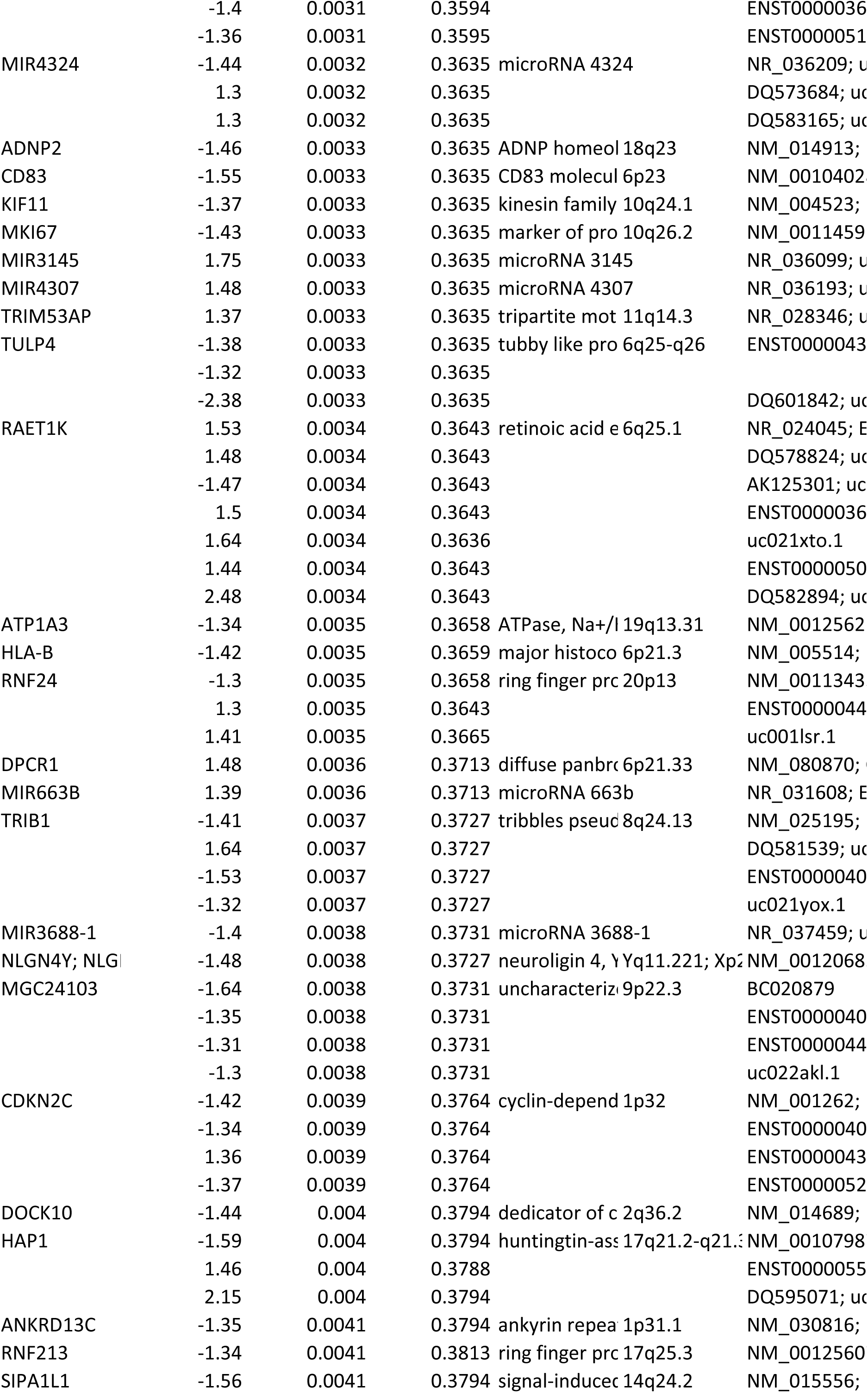
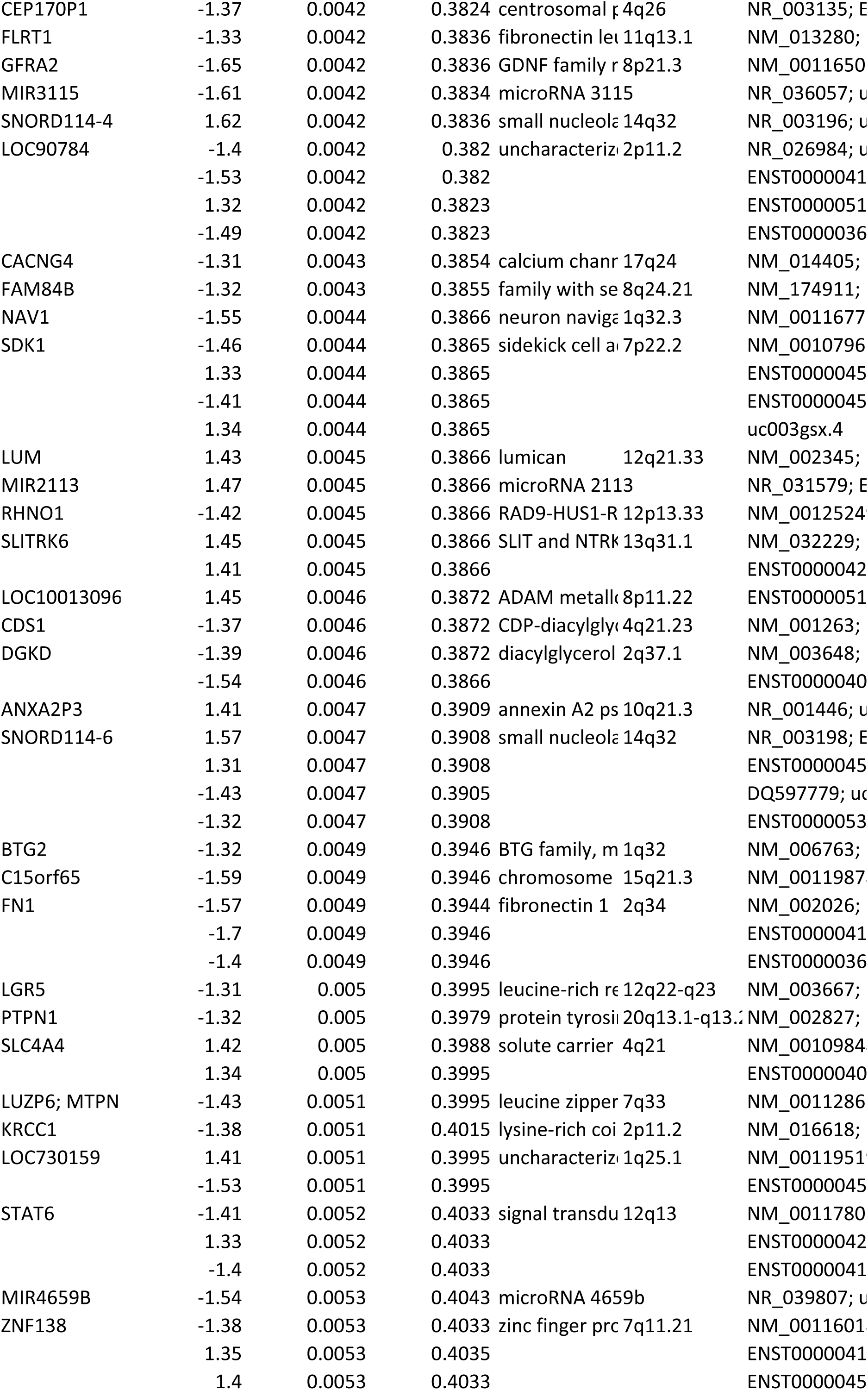
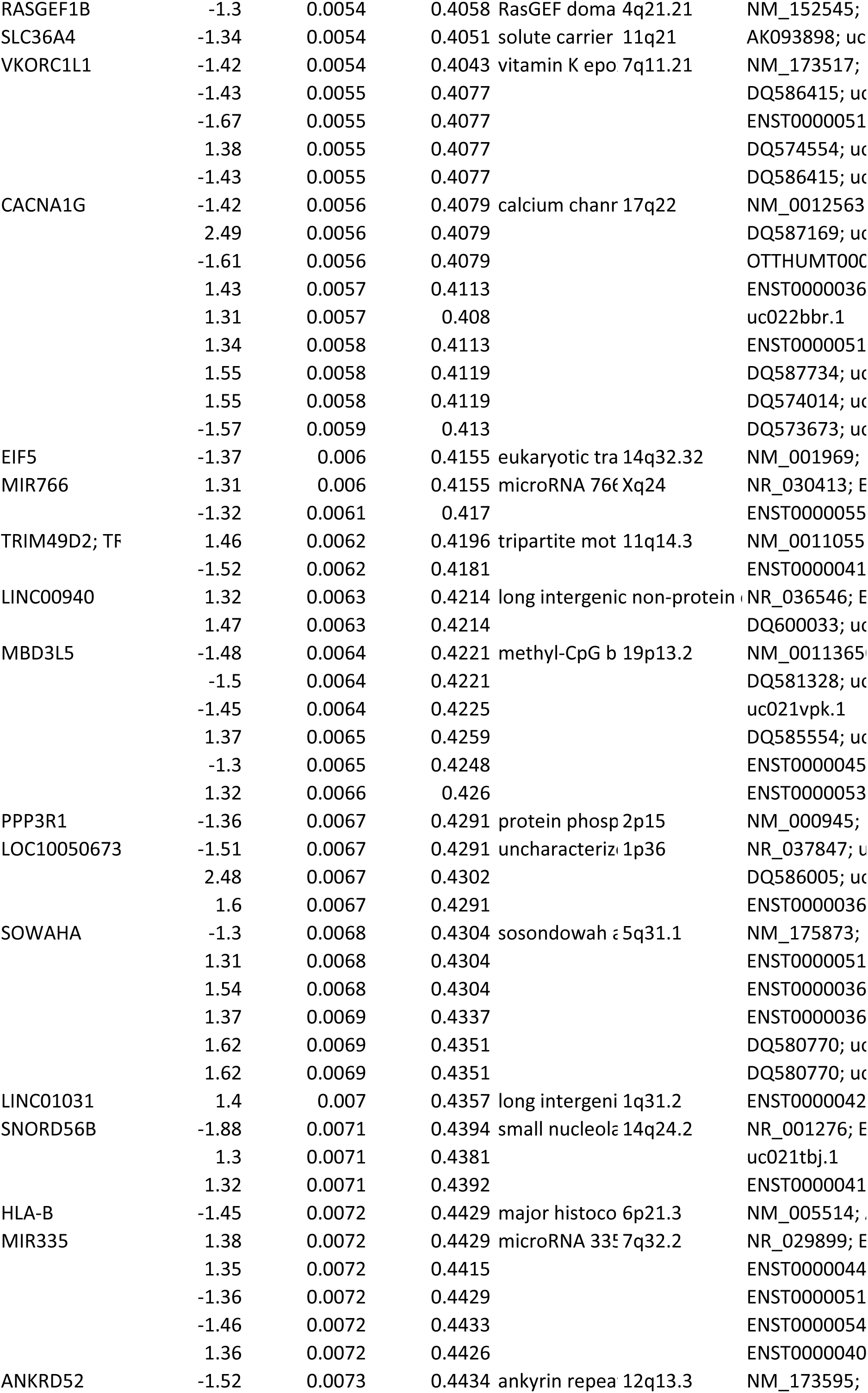
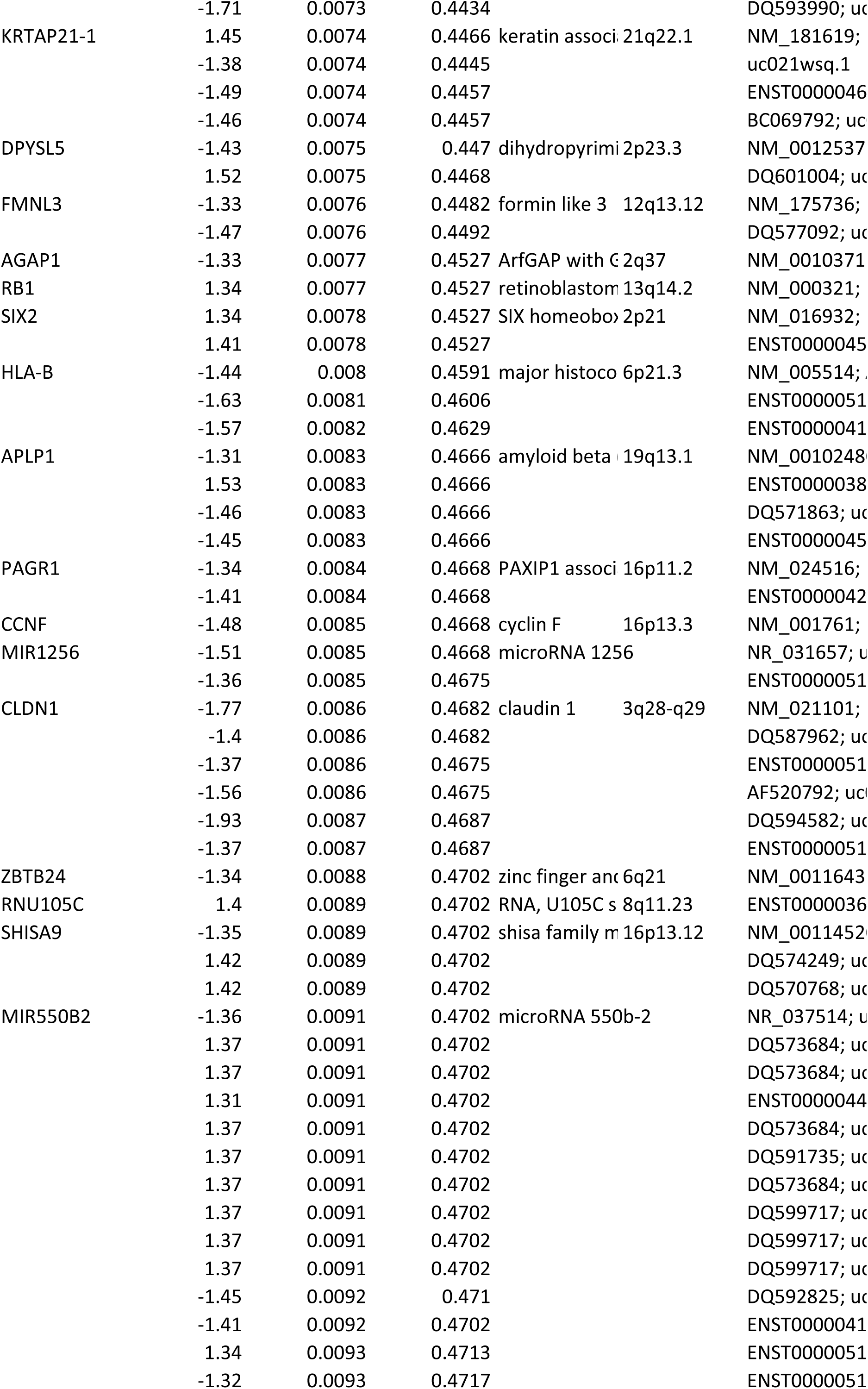
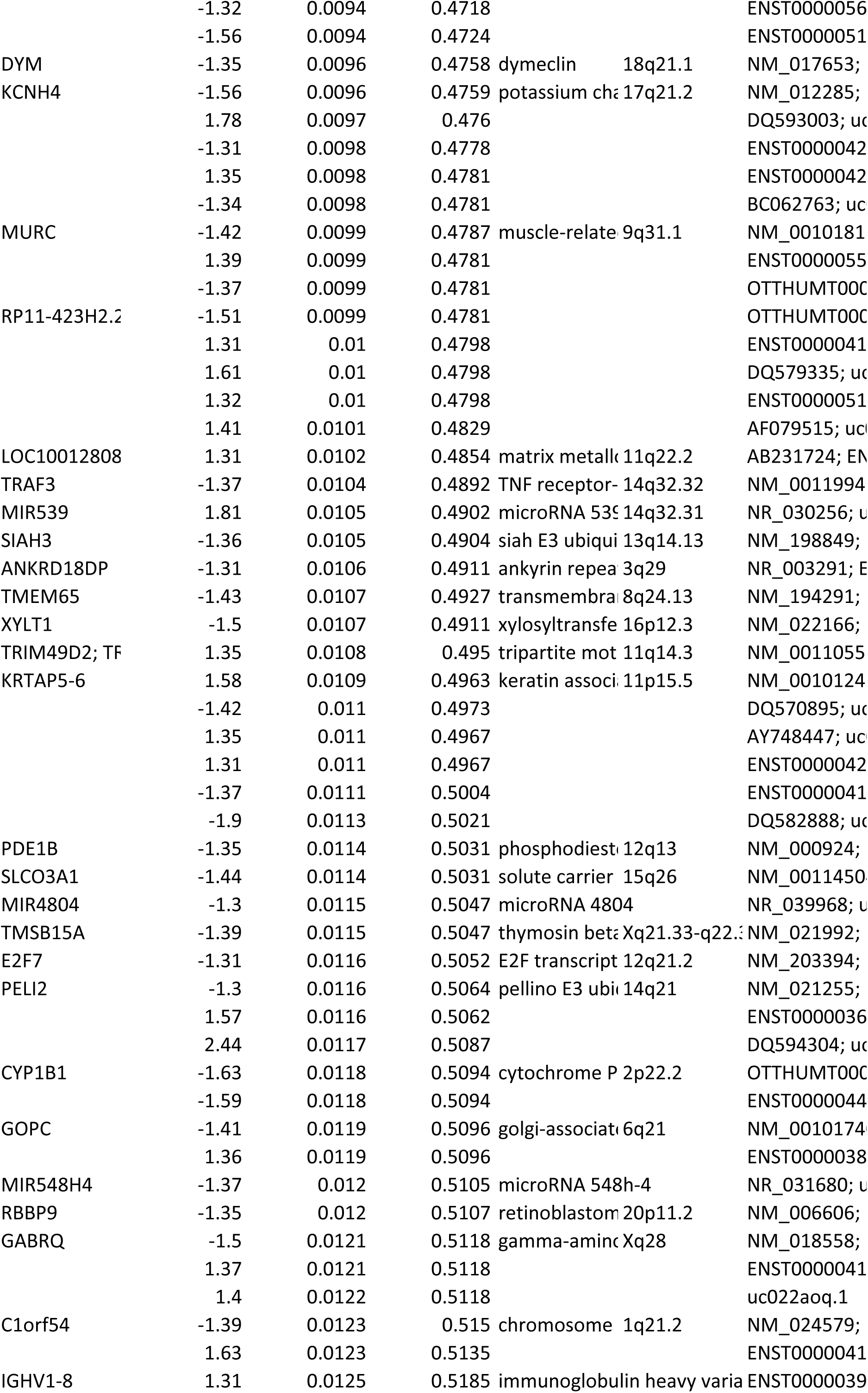
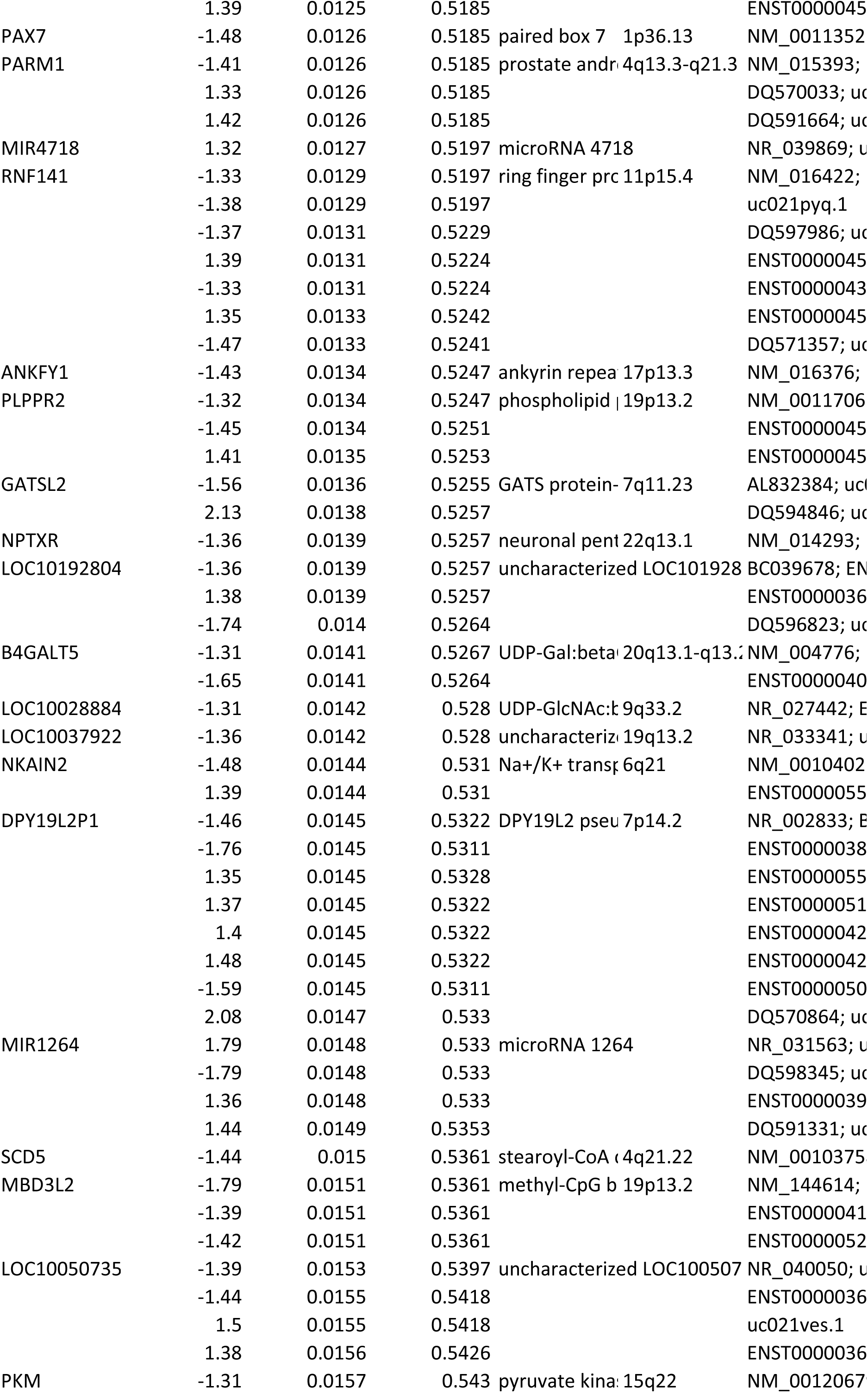
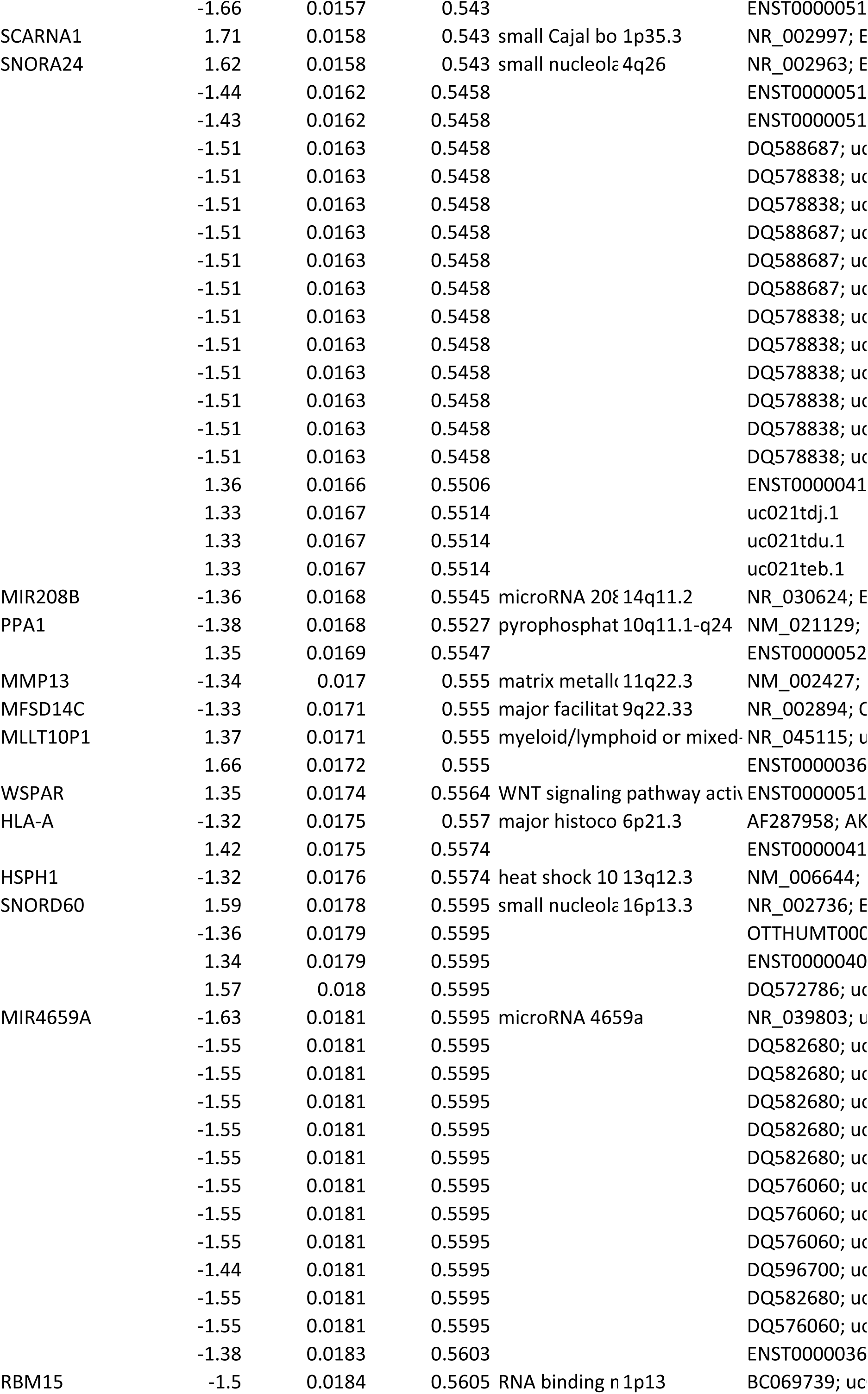
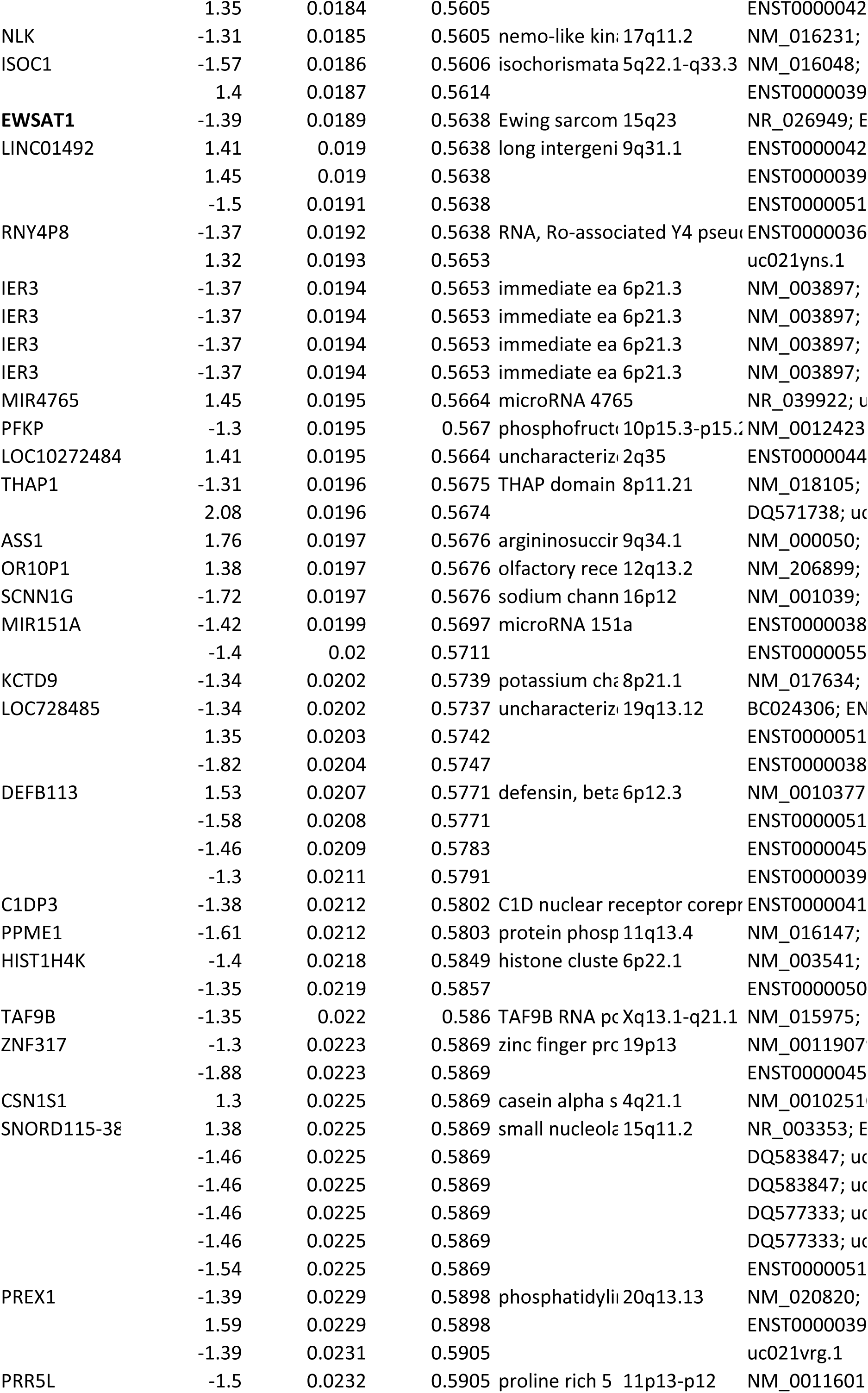
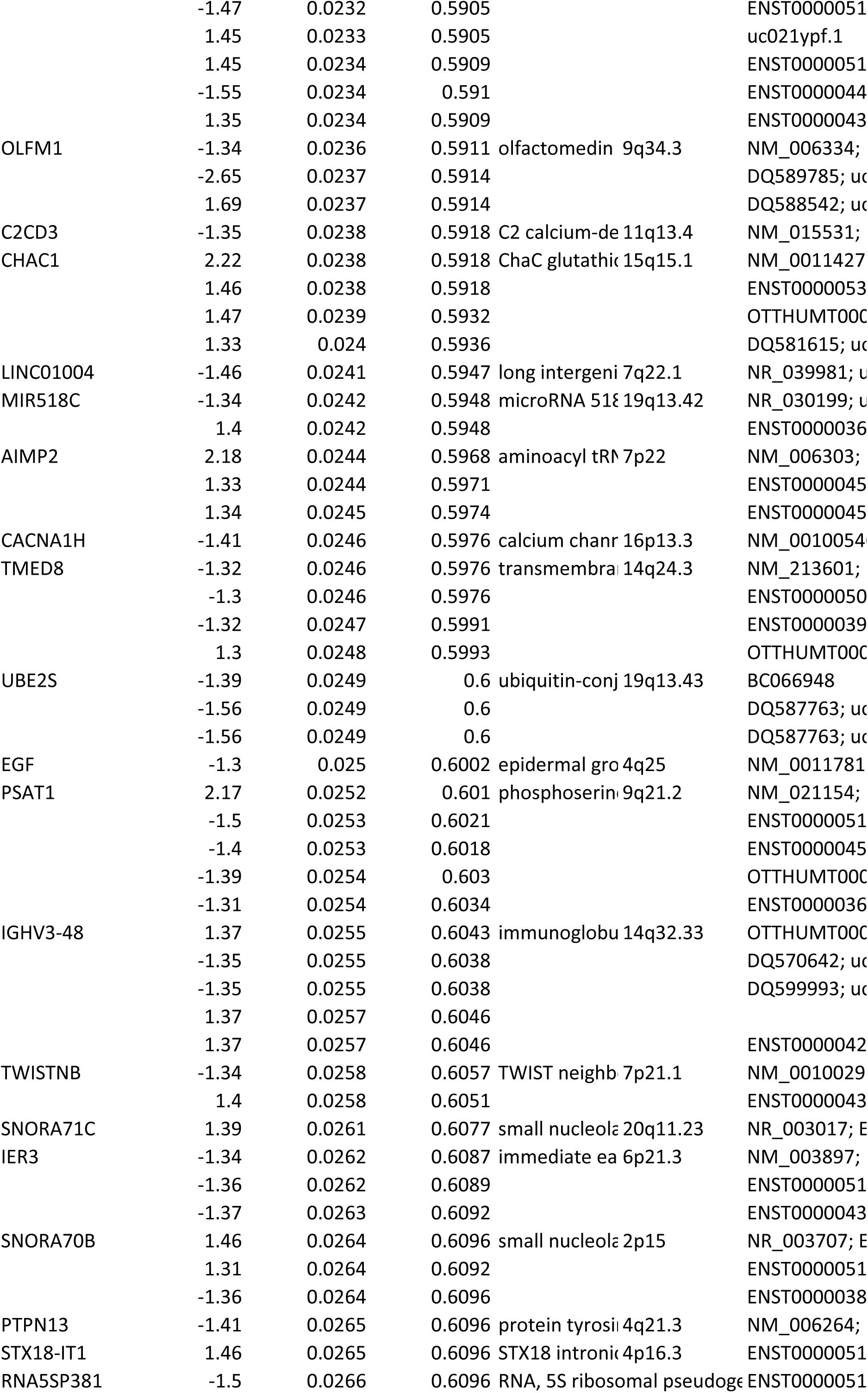
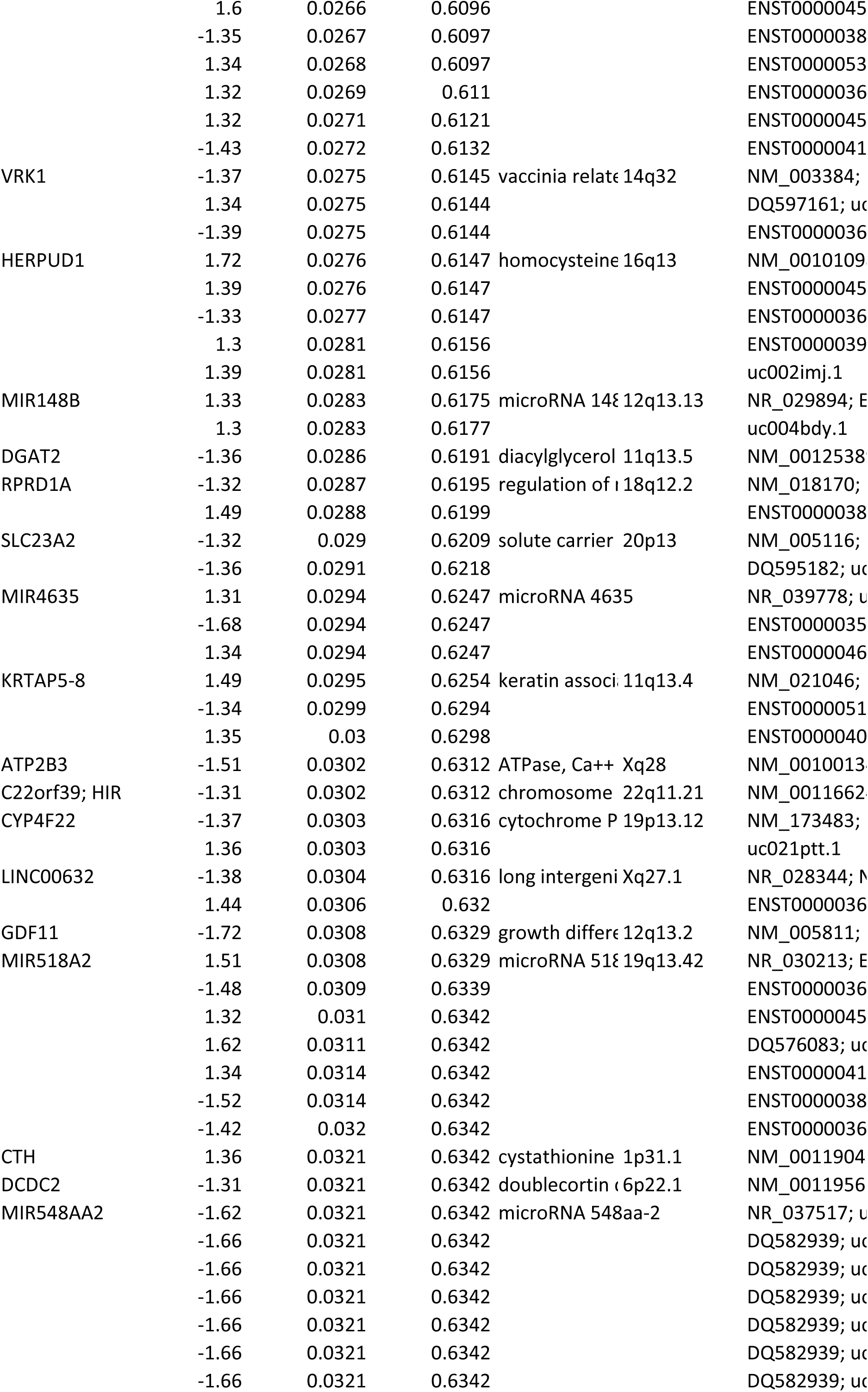
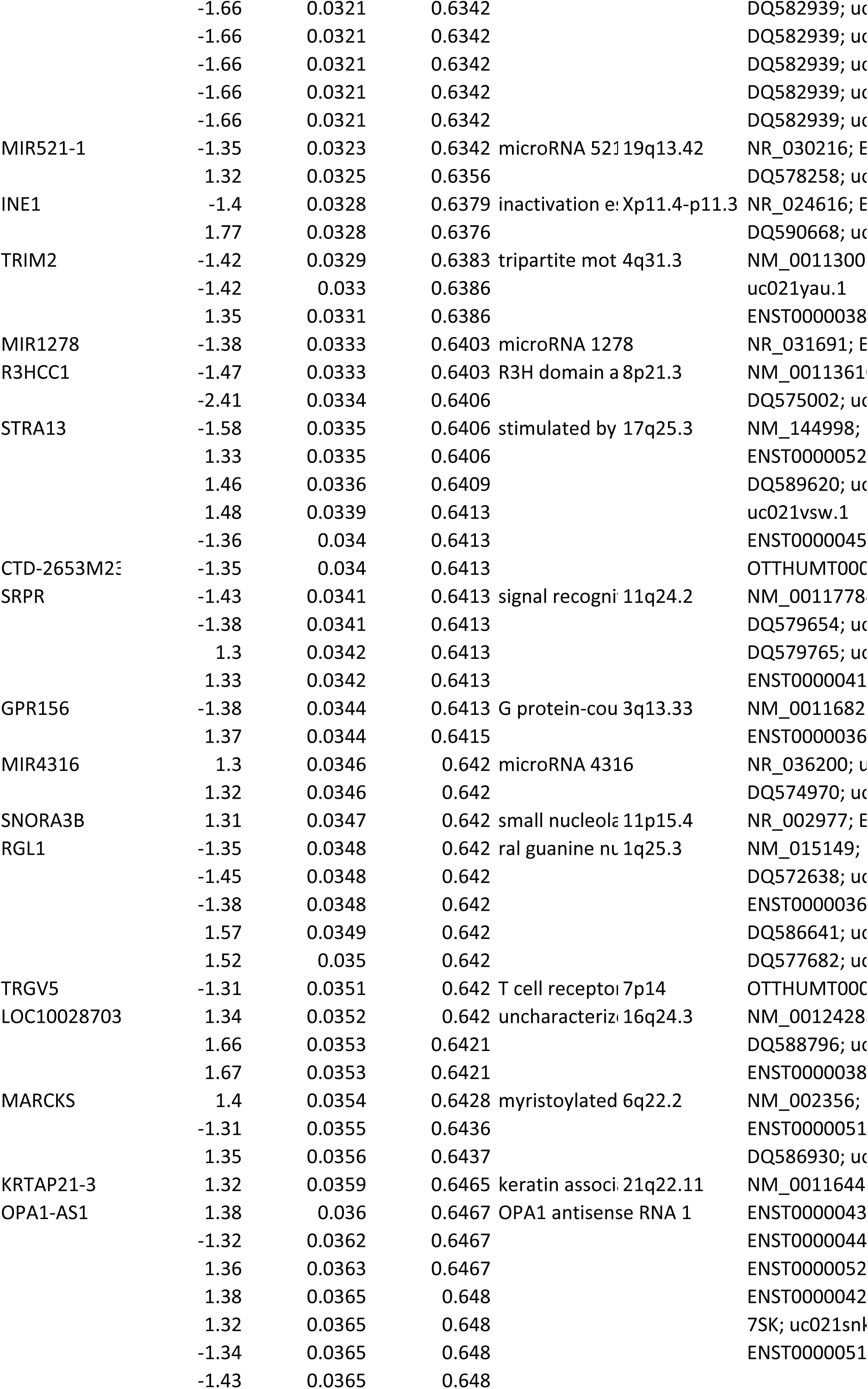
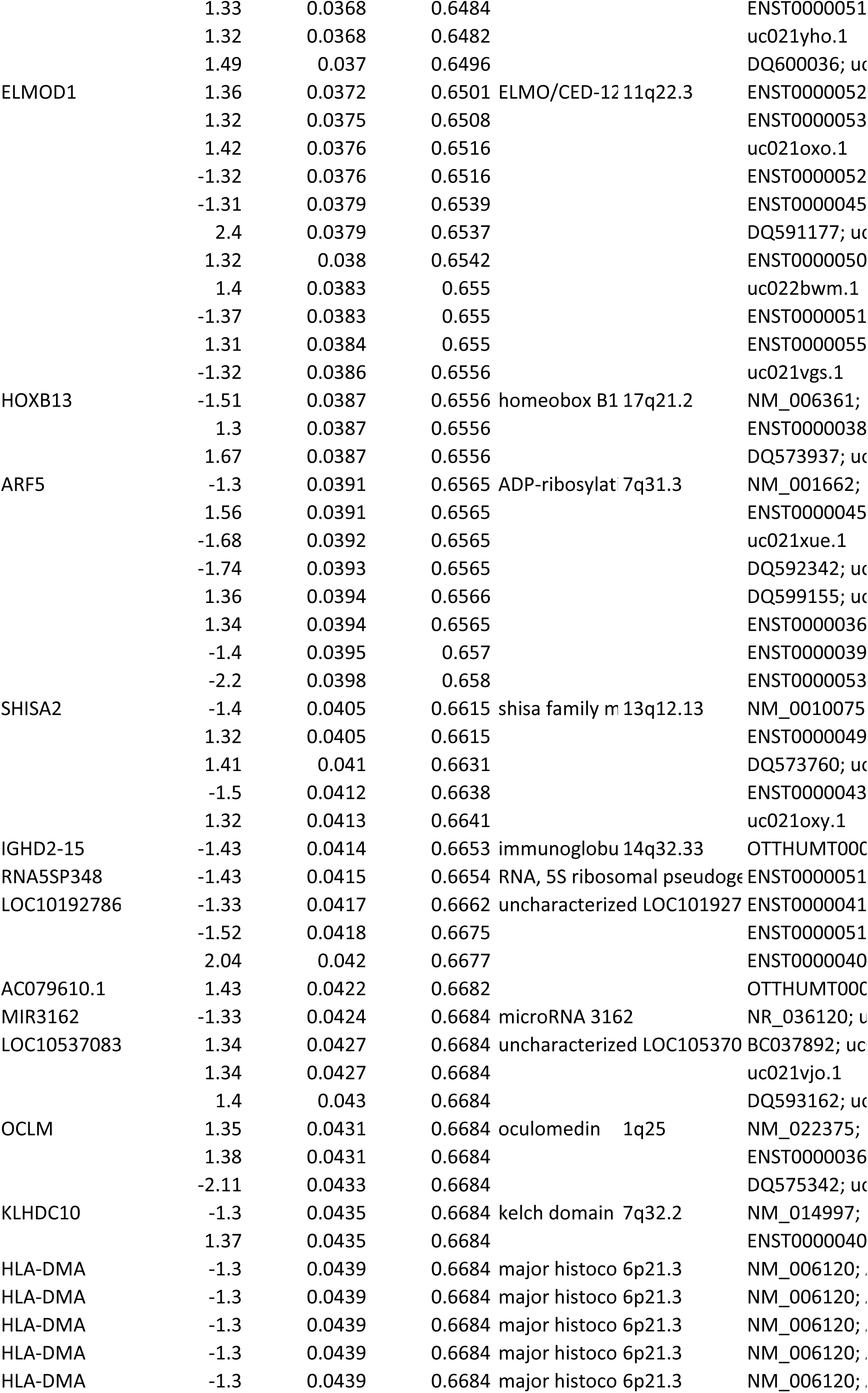
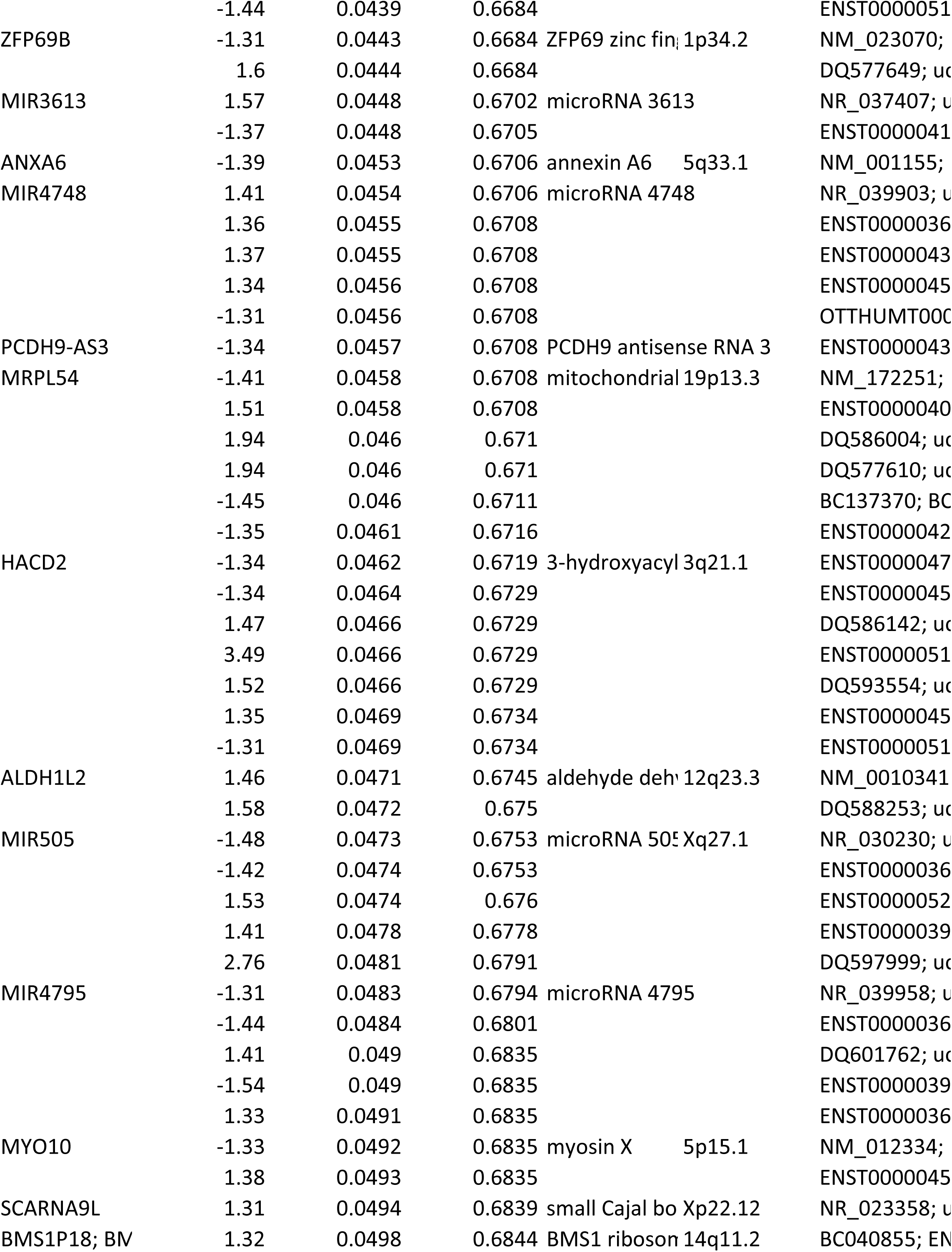
Genes modulated by YAP1/TAZ (siControl vs siYAP1/TAZ)

